# The urban tree of life: synthesizing relationships between body size and urban affinity

**DOI:** 10.1101/2025.09.15.676216

**Authors:** Corey T. Callaghan, Diana E. Bowler, Vaughn Shirey, Brittany M. Mason, Laura H. Antão, Ingmar Staude, John H. Wilshire, Thomas Merckx

## Abstract

Urbanization is a major global driver of biodiversity change, with species responses to urban settings ranging from avoidance to exploitation. To better understand these responses, we conducted a global analysis of urban relative affinity inferred from occurrence data across more than 30,000 animal and plant species. Our synthesis showed a consistent pattern across taxa and biogeographic regions: many species are urban avoiders, while few thrive as urban exploiters—a pattern we coin “Species Urbanness Distribution”. We then assessed whether body size, an integrative ecological trait fundamental to space use, mobility, metabolism, and environmental sensitivity, showed consistent associations with urban affinity among species and across 371 taxonomic families. Analyses were conducted at the interspecific level and focused primarily on variation among taxonomic families (with an accompanying application to view results available here: https://globalecologyresearchgroup.github.io/body_size_results_visualization/). Larger body sizes were generally associated with greater urban affinity in plants compared to animals, though these size-affinity relationships showed considerable variability among families. Our findings highlight the heterogeneous relationship between body size and urban affinity across the tree of life, underscoring the importance of tailored strategies to support urban biodiversity. This research advances ecological understanding of urban filtering and provides a framework for guiding biodiversity-sensitive urban planning amid accelerating global urbanization.

## Main

Cities represent novel anthropogenic environments, leading to significant mismatches with the evolutionary history of most organisms^*1*^. As urban areas are set to expand two- to sixfold over the 21^st^ century^*2*^, urbanization poses a substantial and accelerating threat to global biodiversity^*3*^. Yet species vary widely in their responses: some are completely extirpated (‘urban avoiders’), others persist through behavioral or ecological acclimation (‘urban adapters’) and/or rapid adaptive evolution to cities^*4,5*^, and some even thrive in urban environments^*6*^ (‘urban exploiters’). Quantifying these divergent responses is key for understanding how urbanization will continue to shape global biodiversity and urban ecosystem functioning (e.g., reshaping of food webs^*7,8*^). However, urban ecology has typically relied on small-scale studies, often focused on relatively few species and/or particular taxonomic groups such as birds^*9*^ or mammals^*10*^. Thus, whether general patterns of urban preference or avoidance emerge across species within and across regional pools remains untested.

Species likely do not respond randomly to urbanization as traits may modulate species’ responses. Trait-based approaches are powerful for identifying generalizable patterns across diverse taxa, particularly at broad spatial and phylogenetic scales^*11*^. Although many traits (e.g., diet, reproductive strategy, dispersal ability) may influence species-specific responses to urbanization, body size stands out as both the most widely measured and most taxonomically integrative. It is a key trait of any organism, relating to space use, life-history, and metabolic rate across the tree of life^*12,13,14*^, making it a plausible direct or indirect predictor of species’ responses to key stressor gradients in urban landscapes, including resource fragmentation, heat stress from urban heat islands, and anthropogenic disturbance (Fig. 1).

**Fig. 1:**
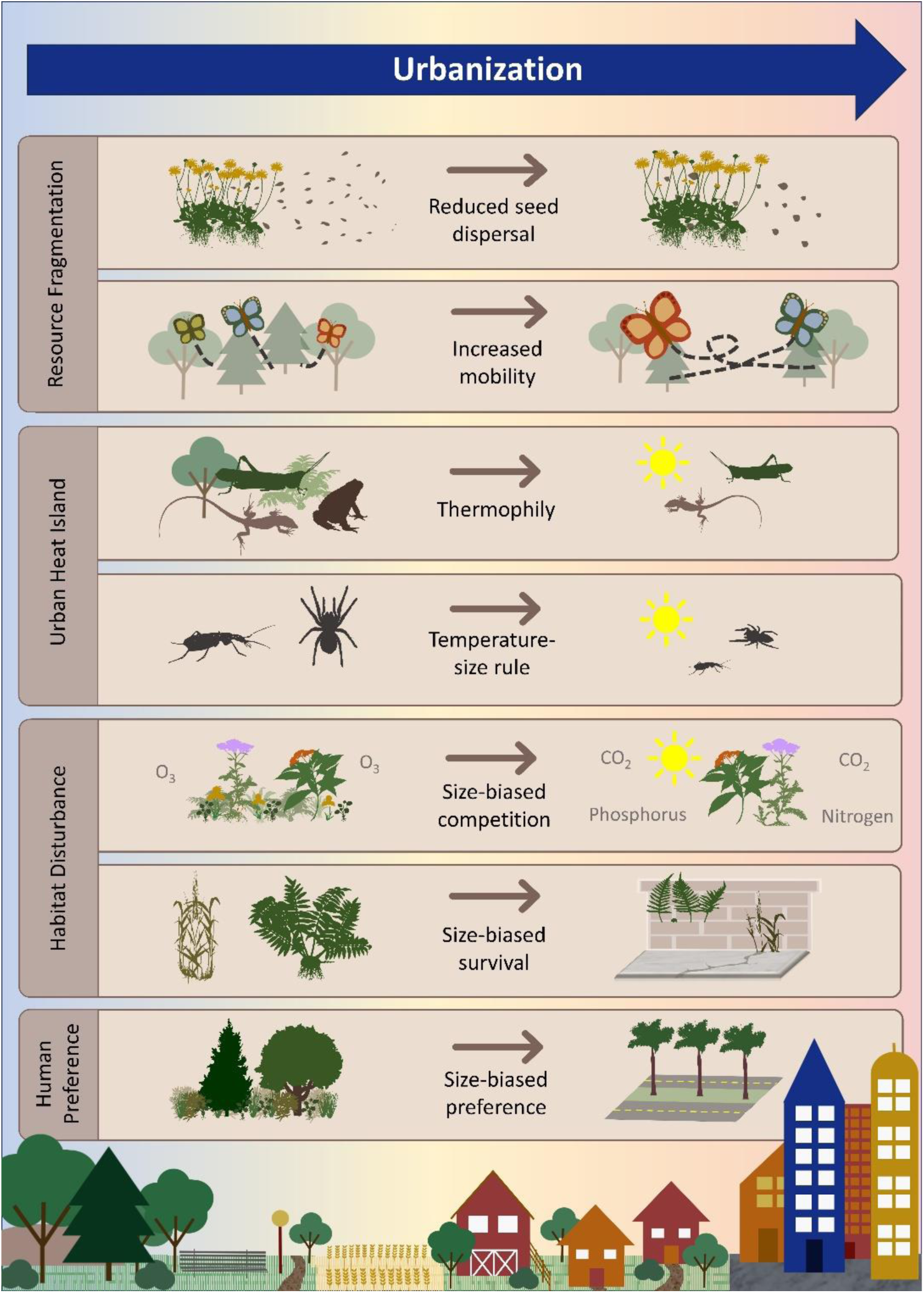
Conceptual framework illustrating hypothesized mechanisms linking urban affinity to interspecific body-size shifts. These include dispersal and mobility constraints under habitat fragmentation^*44,45*^, thermophily and the temperature–size rule driven by the urban heat island effect^*15,30*^, size-biased competition and survival^*94,95*^, and size-biased human preferences^*64*^. Urban fragmentation of habitat resources can select for increased mobility (e.g., larger butterflies) or reduced mobility (e.g., larger seeds) depending on isolation severity. Elevated urban temperatures favor thermophily, which often negatively correlates with size as it affects the heat balance via thermal inertia. Similarly, these higher temperatures generally favor smaller-bodied adult ectotherms because they accelerate development and reduce time available for growth (i.e., temperature-size rule). In plants, the increased CO₂ and nutrient availability associated with anthropogenic environments—due to heating- and traffic-related CO2 emissions and eutrophication—provides a competitive advantage to larger plant species, and human preferences too may favor larger species (e.g., tree-lined streets), whereas smaller species may be advantaged in colonizing built infrastructure.

Body size correlates with urban mobility, but in complex ways^*15*^. For instance, while higher mobility allows some animal species to better cope with the fragmented nature of urban habitat resources, reduced mobility might be advantageous for exploiting localized resources and avoiding risks associated with the urban matrix^*16,17*^. In plants, body size—often indexed by height—correlates with competitive ability, light acquisition, and reproductive strategy, with larger species generally having greater resource needs but potentially higher resilience to urban stressors^*18*^. Notably, the urban heat island effect may drive shifts to smaller-bodied animal species due to elevated metabolic costs^*14,15*^, echoing similar patterns observed under global warming^*19,20*^. However, these shifts toward smaller size can be overruled by requirements for increased urban mobility in taxa where mobility increases with body size^*15*^. Overall, despite body size’s fundamental role in modulating species responses to their environment, a systematic, cross-taxon assessment of how urbanization filters body size distributions is lacking.

Here, we present a global synthesis of urban affinity across more than 30,000 animal and plant species, using occurrence records from the Global Biodiversity Information Facility (GBIF) and remotely sensed night-time light intensity as a proxy for urbanization. We define a continuous metric of regional urban affinity calculated as the realized spatially-explicit urban affinity per species in each subrealm. This metric measuring a species’ relative affinity to urban areas, with negative values indicating avoidance and positive values indicating preference^*21*^. To account for regional variation in both species pools and in species’ urban affinity across their range^*22,23*^, we calculated urban affinity within 52 geographic subrealms (Fig. S1), representing broadly coherent species pools and ecological contexts. Our dataset spans 47 taxonomic classes—from flowering plants (8,980 species) to insects (8,674 species) to mammals (648 species)—and represents a uniquely taxonomically broad assessment. We first introduce the concept of “Species Urbanness Distribution” (SUD) to characterize the composition of a regional community in terms of its species’ urban affinities. We then use Bayesian hierarchical modeling to test whether body size predicts urban affinity across 17,722 species from 371 families, across multiple subrealms. These analyses allow us to quantify both consistency and heterogeneity of body size effects on urban affinity across multiple taxonomic levels, and across regions. This synthesis addresses two core questions: (1) What is the shape of species’ urban affinity distributions within regional communities? (2) Does body size consistently correlate with species’ urban affinities across taxonomic groups and biogeographic contexts? Our aim is to identify broad, cross-taxonomic patterns in species’ urban affinity at a global scale, rather than to resolve the specific causal mechanisms driving urban success or failure within individual taxa or cities.

## Results and Discussion

### Species Urbanness Distributions (SUDs)

We identified a consistent pattern in the extent to which regional communities—defined as all species within a given biogeographic subrealm—consist of urban avoiders and exploiters. We term this the Species Urbanness Distribution (SUD), which captures the full distribution of urban affinity values across species in a community. SUDs exhibited a characteristic shape: a pronounced peak at negative values and a long right-skewed tail toward high positive values, indicating that most species are ‘urban avoiders’, while only a few are ‘urban exploiters’. These patterns in central tendency were broadly consistent across subrealms and taxonomic levels, although distributional shapes varied among higher taxonomic groups (Fig. 2). Conceptually, SUDs parallel the well-known Species Abundance Distributions^*24*^ (SADs), which describe the general law of communities being composed of many rare and few common species^*24,25*^. Similarly, much like the skewed distributions observed in SADs^*24,26*^, the skewed shape of SUDs indicates that while many species exhibit some degree of urban affinity, a relatively small subset of species attain high levels of urban affinity and dominate urban environments. While previous studies have reported a similar pattern at smaller spatial scales^*27*^, our synthesis demonstrates this applies across the globe and taxa. To evaluate this more formally, we compared distributions across subrealms for groups with the largest sample sizes and found that while distributional shapes varied among higher taxa, median values and overall spread were broadly similar within comparable taxonomic levels (Figs. S2–S4).

**Fig. 2:**
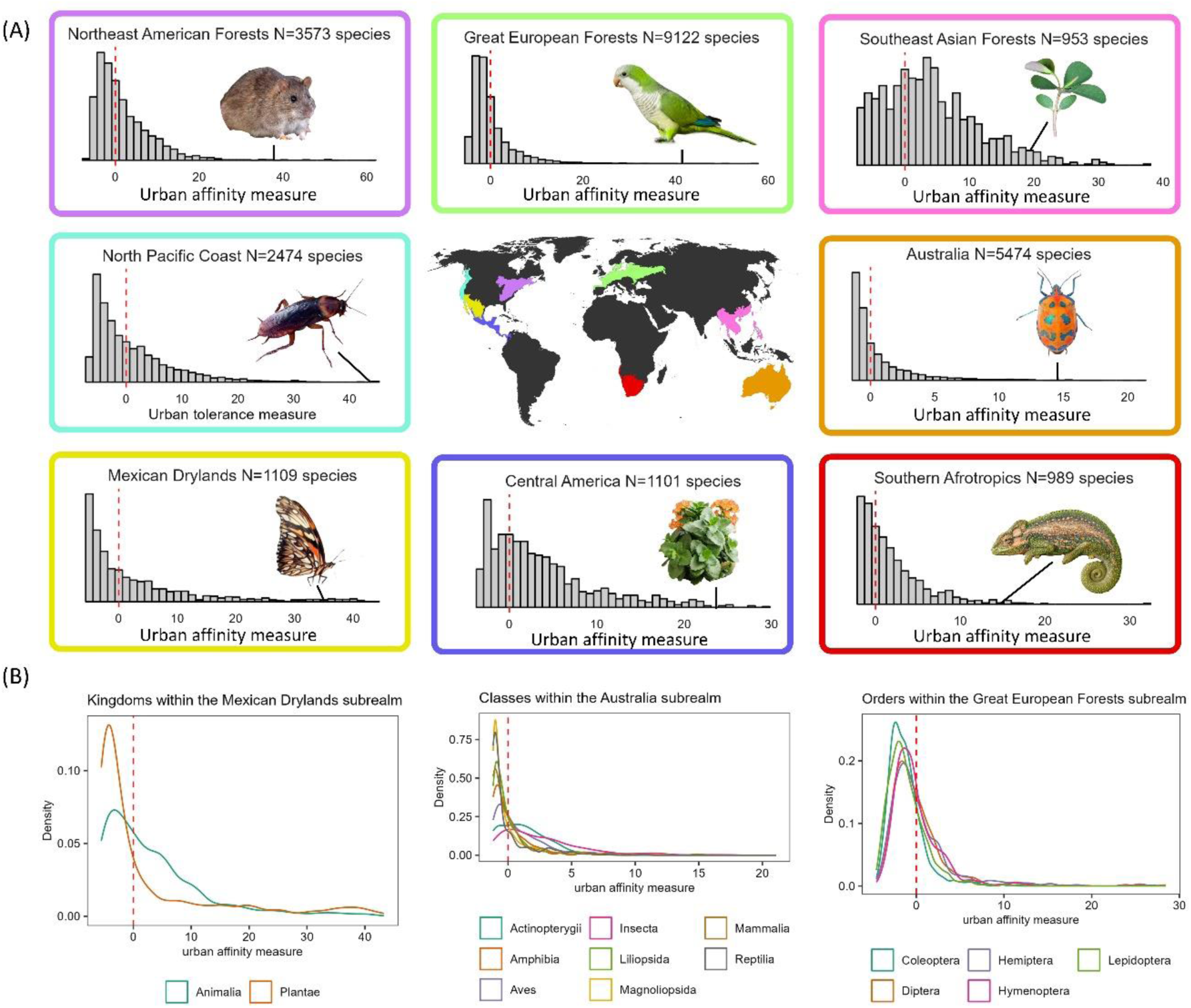
Species urbanness distributions (SUDs) exemplified for eight subrealms. Plotted are all species per subrealm (**A**), with the images highlighting an example ‘hyper-exploiter’ species from each of these subrealms (i.e., with a high urban affinity score). The x-axis shows the urban affinity measure whereas the y-axis is the number of species within that bin. There were consistent patterns for kingdoms, classes, and orders (**B**) as shown by similar central tendencies despite variation in distributional shape. The vertical dashed line represents where species are neutral towards urbanization. Photos acquired from iNaturalist CC BY-NC and background was removed by authors: Brown rat (© Ouwesok), Monk parakeet (© Juan Emilio), Cape dwarf chameleon (© Berkeley Lumb), American cockroach (© Len Worthington), Flaming kay (© Lyubo Gadzhev), Juno silverspot (© Rigoberto Ramírez Cortés), Hibiscus harlequin bug (© Sam Fraser-Smith), and Mascarene island leaf-flower (© Douglas Goldman).

The skewed shape of SUDs suggests that traits enabling species to tolerate urban environments are unevenly expressed, given that only a handful of species show extreme urban affinity values, but our results suggest this is geographically widespread across taxa. These traits may be partly found in human-commensal species, which are found globally and often include invasive non-native species^*28*^. Such traits “pre-adapted” to urban conditions allow for some species to not only persist but thrive in urban environments where most species cannot. Framing these patterns through the lens of exaptation may be particularly useful, as traits that evolved under non-urban selective pressures may incidentally confer advantages in urban environments without having arisen in response to urbanization per se (sensu Lambert et al.^*4*^). We therefore speculate that the skewed shape of SUDs may reflect the uneven distribution of exaptive traits across species pools, rather than widespread adaptive evolution to urban conditions.

Consistent with this interpretation, if exaptive traits that facilitate urban persistence are unevenly distributed across species pools, most species would be expected to exhibit avoidance rather than affinity toward urban environments. Indeed, we found that the median urban affinity is most often below one, indicating widespread avoidance among species (Fig. 2, S5). This is expected, given that urbanization typically involves severe loss and fragmentation of habitat resources for most organisms^*1*^, which negatively impacts numerous taxa, including birds^*29*^, beetles^*30*^, and moths^*31*^. Additionally, the urban heat island—typical of most cities^*32*^—imposes thermal stress, especially during extreme heat events^*33*^, which may drive community shifts toward heat-tolerant species, as observed in ants^*34*^, bees^*35*^, and plants^*36*^. Yet, urban environments also frequently contain green infrastructure and remnants of natural habitats that can act as sanctuaries for rare and endangered species^*37*^, and cities can harbor a large part of the regional diversity^*38*^. But these green infrastructure spaces are often also dominated by widespread generalist species. The multitude of environmental stressors, combined with competition dynamics between urban-tolerant and intolerant species, likely influence the shape of SUDs (Fig. 2). Future ecological research should aim to disentangle these contrasting drivers across taxa and regions to better understand the ecological filters at play.

### The relationship between body size and urban affinity

To test whether there is a consistent pattern of body size filtering associated with urban affinity across the tree of life, we integrated our species-level urban affinity values with body size measures collated from the literature. We performed this analysis for 17,722 species spanning both plants (i.e., plant height) and animals (e.g., body length, mass, wingspan). We first evaluated the overall relationship between body size and urban affinity across all taxa, fitting two models (see Methods for details). We found that the effect of body size on urban affinity is stronger in plants than in animals (0.64 vs. 0.21), although uncertainty was substantial in both groups, with zero-overlapping 95% credible intervals (plants: -0.68 to 1.84; animals: -0.25 to 0.66). Variation in urban affinity was most pronounced at the family level (sd = 2.43), followed by class and order level (sd = 2.34 and 1.75, respectively). Similarly, the effect of body size varied most among families (sd = 1.24), compared to orders (sd = 0.73) and classes (sd = 0.27), suggesting that body size predicts urban affinity most consistently within the finer family-level taxonomic scale, rather than at higher scales where other factors may obscure relationships. Based on these patterns, we focused subsequent analyses at the family level, fitting Bayesian models for families with data on at least ten species to independently explore the relationships between urban affinity and body size. Because body size covaries with multiple ecological traits (e.g., dispersal ability and metabolic rate), we focused on family-level analyses to capture shared ecological strategies while still allowing sufficient variation among species to detect trait–environment relationships^*39*^.

Across 371 families (93 plant and 278 animal families), we found no universal relationship between body size and urban affinity; instead, patterns were heterogeneous (Fig. 3). Using 80%, 90%, and 95% credible intervals to infer weak, moderate, and strong effects, respectively, we found evidence of body size filtering in 83 families (29 strong, 19 moderate, and 35 weak). These included 48 animal families (17% of animal families) and 35 plant families (38% of plant families), indicating that for most families, body size does not consistently predict urban affinity. Nonetheless, among plant families, the effect of body size on urban affinity was predominantly (N=77; 83%) positive, with 23 families showing strong or moderate effects. In contrast, animal families (N=278) exhibited more variable responses: 58% showed positive effects and 42% negative, with only a subset showing strong or moderate effects (positive: 18 families; negative: 6 families). Notable families with the strongest positive associations included Salicaceae (willows) and Columbidae (doves and pigeons), where larger species are more urban-tolerant. In contrast, Dipsadidae (snakes) and Accipitridae (birds of prey) showed the strongest negative associations. Further discussion of notable order-level effects is provided in the Supporting Information. While the overall patterns for family-level effects are visible in Fig. 3, we also developed an interactive online visualization of family-level effect sizes, available here.

**Fig. 3:**
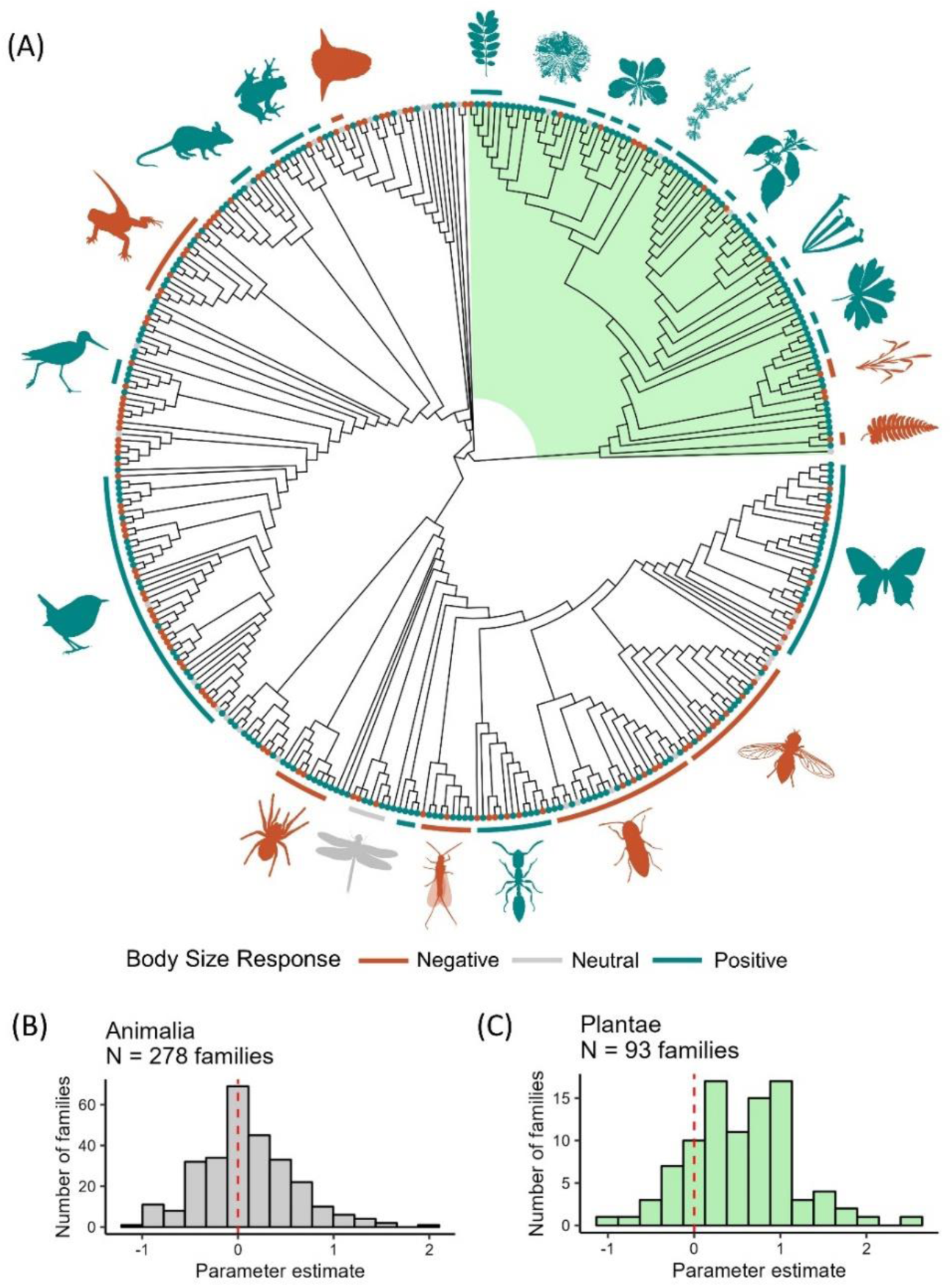
Effect sizes between body size and urban affinity across the tree of life and individual effect sizes for animals and plants families. (**A**) Effect sizes of the relationship between body size and urban affinity for the 371 families included in our analysis, plotted along a phylogenetic tree of life (see Methods); Plantae are highlighted and shaded in green. Colors indicate the direction of the effect: orange indicates negative, petrol indicates positive, grey indicates neutral (i.e. any effect sizes between -0.05 and 0.05). (**B**) and (**C**) histograms of individual effect sizes for each family, for animals (B) and plants (C). Orders are shown along the outside edge of the phylogenetic tree, each with a bar and icon, for any order with more than 3 families. An interactive version for full exploration of our results at both family and order level is available here.

The variability in both the direction and strength of urban body size filtering across taxa likely reflects the diversity of urban stressors and the contrasting taxon-specific pathways through which body size mediates sensitivity to them. In raptors, for example, the negative association between body size and urban affinity—consistent with previous findings^*40,41*^—likely stems from large species’ need for extensive hunting territories, whose availability and access are constrained by urbanization-driven habitat loss and fragmentation. Similarly, the strong negative effect observed in Coleoptera supports the hypothesis that the urban heat island drives shifts to smaller body sizes in urban-tolerant ectotherms, consistent with Atkinson’s temperature-size rule^*42*^. Larger beetles may also be more vulnerable to urban disturbance due to lower reproductive output and reduced dispersal capacity^*43*^. In contrast, urban communities of Lepidoptera (moths and butterflies) typically shift toward larger—and hence more mobile—species than rural communities^*15*^, likely because greater mobility facilitates persistence in typically fragmented urban landscapes^*44*^.

Our results are broadly consistent with prior taxon-specific trait-based studies (e.g., Hahs et al.^*17*^), but also highlight that relationships between body size and urbanization vary across taxa and analytical frameworks. For example, global syntheses and regional studies have reported positive, negative, or null size–urbanization relationships depending on clade and spatial scale. A recent global analysis that compiled empirical occurrence data for multiple terrestrial faunal taxa across cities worldwide reported broadly similar body-size responses to urbanization^*17*^. For four of the five groups that overlap with our analysis—amphibians, bats, bees, and birds—the direction of the body-size relationship with urbanization was consistent between studies. The only exception was carabid beetles, which tended to be smaller-bodied in highly urbanized environments in that analysis, whereas we detected no significant size effect for this family. Studies on birds, for example, have found mixed results, including positive associations to urbanization in some regional assemblages^*45*^, no global relationship in others^*46*^ or an overall negative relationship globally^*23*^, and negative relationships in particular clades such as raptors^*40*^. Such discrepancies likely arise because different studies quantify urbanization differently, focus on different spatial grains, or analyze different components of species responses (e.g., presence–absence, abundance, or occurrence distributions). Additionally, a study on multiple taxa including butterflies and moths found a positive relationship in butterfly and moth community-weighed mean body size with increases in urbanization level, similar to our findings^*31*^. Researchers have also found that smaller-bodied dung-associated beetles potentially benefit from urban environments, which is similar to the negative association we found between urbanization and body size in beetles^*47*^. Our approach complements these studies by estimating occurrence-based urban associations across thousands of taxa simultaneously, allowing comparison of how consistently body size predicts urban affinity across taxonomic groupings rather than within a single lineage. In this sense, variation among published results does not contradict our findings but instead reinforces the conclusion that body size is a context-dependent filter whose direction and strength depend on ecological setting, taxonomic scope, and the urbanization metric used.

Order-level analyses (see Supplementary Text and our interactive figure here) offer further insight into the mechanisms shaping body size responses to urbanization. In plants, we consistently observed a positive association between body size and urban affinity, except for grasses and ferns. Taller species may gain a competitive edge in urban settings^*48*^, by exploiting higher temperatures, increased CO_2_ and nitrogen availability^*49*^, and typically reduced O_3_ concentration^*50*^. In contrast, smaller species of grasses and ferns were favored by urbanization in our study, likely stemming from high disturbance regimes in urban grasslands and ruderal environments. Among ectotherm animals—including Diptera, Coleoptera, Hemiptera, and Squamata—the general expectation is a shift toward smaller species, consistent with metabolic constraints imposed by the urban heat island (Fig. 1, Fig. 3). However, exceptions occur when urban fragmentation requires greater mobility, favoring larger and more mobile species^*15*^. Urban spiders also exhibit body size reductions, driven by both elevated temperatures and reduced prey size^*51*^. For endotherms such as rodents and birds, we typically anticipate a shift toward larger species because of increased habitat fragmentation, especially when body size correlates positively with mobility. While this pattern held for most bird families, notable exceptions—like Piciformes and Accipitridae—demonstrate negative associations, possibly due to constraints related to large home range requirements and prey size^*52,53*^. Future research should further explore such trophic consequences of urban size filtering, as these effects can either align across trophic levels or become mismatched. For instance, urban environments may favor larger predators—such as insectivorous birds and mammals—while simultaneously selecting for smaller insect prey species. Such mismatches can lead to nutritional stress, reduced reproductive success, and ultimately a more severe homogenization of urban predator guilds^*54,55*^. Another ecological consequence of size mismatches may occur when urban pollinator assemblages become disproportionally large or small relative to the floral traits of native plants, potentially reducing pollination efficienc^*56*^—although rapid evolutionary shifts in floral morphology could help mitigate these effects^*57*^. Because our synthesis is correlative and macroecological in nature, the mechanisms discussed above are best viewed as hypotheses that can be evaluated through future work combining experimental, trait-based, and longitudinal data.

Ultimately, the heterogeneous and sometimes weak relationships between body size and urban affinity suggests that body size alone cannot explain the emergence of extreme urban exploiters and the skewed shape of SUDs. Focusing on body size as a focal trait necessarily represents a simplification of the multidimensional processes underlying species’ responses to urbanization, driven in part by data availability when conducting a taxonomically-broad synthesis. Instead, urban affinity likely depends on multivariate trait combinations^*17,58*^ that vary among taxa^*59*^ and ecological contexts^*60*^. Traits that are likely to correlate with urban affinity include dispersal capacity, behavioral flexibility, diet breadth, reproductive strategy, thermoregulatory ability, and, in plants, life history traits such as growth form, clonality, phenology, and seed size. The diversity of trait pathways through which species may persist or thrive in urban environments is consistent with the pronounced taxonomic heterogeneity we observe and helps explain why body size alone does not yield a universal pattern.

## Current limitations and future directions

Our analysis, which incorporates over 17,000 species, represents the most extensive assessment to date of how body size relates to urban affinity across the tree of life. Such scale was enabled by the growing mobilization of biodiversity data through GBIF, particularly from citizen science platforms like iNaturalist^*61*^. Still, our analysis largely focused on common species, as we applied a cutoff of at least 100 observations per species per subrealm. As a result, rare or less frequently observed species are underrepresented, and their responses to urbanization remain an important avenue for future study. This is especially important given the growing recognition of cities as sanctuaries for various species^*37*^. Nevertheless, our findings highlight the relevance of common species to understanding macro-ecological patterns (sensu^*62*^). We focused here on broad-scale patterns, but future work should further explore the context-dependency of body size filtering—for instance, how it varies with city size, urban green infrastructure configuration, regional climate, and human cultural preferences. For example, one possible explanation for the observed trend of larger, urban-tolerant plants may lie in the widespread use of non-native woody ornamentals, such as alien trees and tall shrubs, which reflect human aesthetic preferences^*63,64*^—suggesting human preferences are an additional filter on plant size. These human-driven preferences may also influence detectability and recording effort, as larger and more conspicuous plant species are more likely to be planted, maintained, and documented in urban environments, and thus be available in GBIF for our analyses. However, we suggest that this is not purely a sampling artifact, but such processes likely interact with ecological filtering to shape the realized size structure of urban plant communities. Similarly, human–wildlife conflict and active management of large-bodied animals in cities may influence which species persist in urban environments, potentially constraining the upper end of the body size distribution. Taken together, these examples illustrate the importance of considering the socio-ecological context of urban species assemblages^*65*^. In parallel, while our study examined interspecific variation, growing evidence also points to important intraspecific responses, where body size shifts within species can modulate urban affinity^*31,66,67*^.

One important limitation of our synthesis is the heterogeneity in how body size is measured across taxa, including differences among mean, maximum, and sex-specific estimates. While our analytical framework explicitly accounts for this variation through transformation, scaling, and hierarchical modeling with random intercepts (see Methods), residual measurement noise may still obscure weak size–urban affinity relationships. This challenge is inherent to large-scale trait syntheses that integrate data from disparate sources, and highlights the need for continued efforts to standardize trait databases and expand the availability of harmonized organismal trait data across the tree of life. Nevertheless, the urban affinity scores presented here will form a valuable foundation for future research and local-scale planning efforts, in particular those that use citizen science to track restoration progress. For example, these affinity scores could inform urban greenspace surveys aimed at calculating integrity indices—providing a repeatable, quantitative tool to assess and monitor the long-term success of urban ecological restoration by measuring the ‘urbanness’ of local species communities in urban greenspace^*45*^. In this context, incorporating taxon-specific body size-urban affinity relationships can enhance conservation outcomes by tailoring strategies to the size dynamics of assemblages. For instance, in assemblages where larger species are disproportionally filtered out (i.e., a negative size-affinity link), efforts should focus on increasing the size and quality of habitat patches and mitigating the urban heat island.

Conversely, in assemblages where smaller, less-mobile species are more vulnerable, improving functional connectivity between habitat patches should be prioritized^*68*^. For butterflies in particular, Pla-Narbona *et al*.^*68*^ advocate for such context-specific management strategies—balancing habitat patch quality and connectivity—to promote more diverse urban butterfly communities.

## Conclusions

Our global analysis of over 30,000 species provides new insight into how diverse taxa respond to urbanization, revealing a consistent skew in urban affinity—characterized by many urban avoiders and few exploiters—across biogeographic regions and taxonomic groups. We introduce the concept of Species Urbanness Distributions (SUDs) as a novel framework to quantify and compare the impact of urbanization on species communities. Much like Species Abundance Distributions (SADs) are essential for ecology and biodiversity research, SUDs offer a generalizable lens through which the community structure and ecological resilience of urban biotas can be assessed. Understanding the processes that generate SUDs, and understanding the ecological impacts of differently shaped SUDs, represent key avenues for future research.

Although body size emerged as a predictor of urban affinity, we found not only substantial heterogeneity across families and orders, but also that body size filtering alone is unlikely to explain the consistently skewed SUD shape. Taken together, these patterns suggest that urban affinity likely emerges from multiple trait combinations rather than a single, universally advantageous trait, and that strong affinity to urban environments is not uniformly expressed across taxa, despite occurring broadly across regions. Nevertheless, trait-based approaches—especially those integrating multiple functional traits—hold strong potential for uncovering the processes driving the diversity of species’ urban responses and for interpreting the shape and skew of SUDs. Moreover, trait-based predictions of species’ vulnerability could be used to formulate effective strategies to promote biodiversity in urban environments^*17*^. Our synthesis complements taxon-specific, presence–absence trait studies by identifying broad, cross-taxonomic patterns that can motivate and contextualize more mechanistic analyses^*17,23*^.

Looking ahead, the continued growth of citizen science data platforms, such as iNaturalist, will play an increasingly critical role in tracking and understanding the mechanisms driving biodiversity responses to urbanization. These data streams will enable broader quantification of species’ urban affinity, including rare and currently underrepresented taxa. Combined with trait-based modeling, this growing dataset could be leveraged to identify urban-vulnerable species and hence guide urban habitat restoration initiatives. In this way, SUDs and trait-informed predictions offer powerful tools for more effective urban strategies to conserve biodiversity on an increasingly urban planet.

## Methods

Our methodological approach can be broken down into three key steps: (1) quantifying species-specific urban affinity scores, stratified by subrealm (i.e., biogeographic region); (2) collating measures of body size for as many species as possible for which we were able to quantify urban affinity; and (3) quantifying the relationship between urban affinity and body size at various taxonomic levels.

### Quantifying urban affinity

Our aim was to derive a continuous, occurrence-based metric that describes how species are distributed along an urbanization gradient within a given biogeographic region. To do this, we followed a three-step procedure (sensu Callaghan et al.^*21,69,70*^). First, we quantified the level of urbanization associated with individual species’ occurrence locations (i.e., coordinates) using remotely sensed night-time lights. Second, we summarized these values at the species level within each subrealm to obtain a species-specific regional urban score. Third, we expressed each species’ urban score relative to the regional background of urbanization level to obtain a subrealm-specific measure of urban affinity, which reflects whether a species tends to occur in more or less urbanized environments than is typical for that region. In the following paragraphs, we walk through each of these steps in detail, including the assumptions, limitations, and intended interpretation of the resulting metrics.

### Assigning urbanization level to species’ occurrences

We downloaded occurrence data from the Global Biodiversity Information Facility (hereafter GBIF) (https://doi.org/10.15468/dl.4dcbgt) on February 4th, 2021, including ∼1.4 billion biodiversity records from >24,000 datasets. GBIF data were filtered to only include observations that were recorded as species, and an additional step was taken to ensure that the listed genus in GBIF matched the genus of the species name, ensuring validity of the taxonomic nomenclature of GBIF. Due to uncertainty in matching observations with remotely-sensed products, any GBIF observation with a coordinate uncertainty > 1 km was removed. This filtering step removed individual observations with high spatial uncertainty, rather than excluding entire datasets or survey types. We only included species from a taxonomic Class (i.e., the rank between Phylum and Order) that had at least 10 species reported to GBIF, focusing on the most common classes for downstream analyses.

We then overlaid GBIF species occurrence observations with a remotely-sensed layer representing a continuous proxy of urbanization—Visible Infrared Imaging Radiometer Suite (VIIRS) night-time light^*71*^. We used this proxy as it is a continuous measure of urbanization commonly used to represent urban extent^*72,73,74*^. Previous work has shown that VIIRS night-time lights is negatively correlated with greenness measured through the Enhanced Vegetation Index (EVI) and positively correlated with human population density^*69,71*^. Although night-time light intensity can vary among cities with similar impervious surface due to differences in land use, infrastructure, and cultural lighting practices, at broad spatial scales it functions as an integrative proxy of urbanization^*75,76,77,78,79,80*^, with localized heterogeneity contributing primarily to additional variance rather than systematic bias. We used Google Earth Engine^*81*^ for our geospatial processing and data were obtained for the VIIRS Stray Light Corrected Nighttime Day/Night Band Composites product, representing monthly composites, (i.e., this dataset in Google Earth Engine: [NOAA/VIIRS/DNB/MONTHLY_V1/VCMSLCFG](https://developers.google.com/earth-engine/datasets/catalog/NOAA_VIIRS_DNB_MONTHLY_V1_VCMSLCFG#description)) with a native resolution of ∼500 m^2^. We took the median of all monthly composites for each pixel (i.e., a single grid cell of the nighttime lights raster representing a fixed ground area) to calculate a pixel-level urbanization value, measured in average radiance, using imagery from January 2015 to January 2021. GBIF data were filtered from January 1st, 2010 to February 4th, 2021. We acknowledge that these two temporal scales do not precisely match, but because urban conversion at the landscape/regional scale happens relatively slowly, we assume that the level of urbanization between 2015 and 2021 accurately captures the relative differences between regions with low and high urban land cover.

To avoid computational overload, we used geohash encoding to assign every record in GBIF to a “pixel” representing the VIIRS average radiance based on its geographic coordinates. We used geohash7 encoding to divide the geographic area into grid cells (referred to as blocks), each representing an approximate spatial area of 150 m^2^. The VIIRS night-time lights data, with a native resolution of ∼500 m^2^, was then matched to these blocks by assigning each geohash7 block the average VIIRS radiance value that intersects it. We do not assume positional accuracy at the scale of the geohash blocks, but geohash encoding was used solely for computational indexing, while the effective spatial resolution of the urbanization metric is that of the VIIRS data (∼500 m). This approach allows us to avoid unnecessary redundancy in the data while maintaining the original VIIRS resolution.

### Accounting for geographic context through subrealm stratification

To account for geographic heterogeneity in both species’ distributions and the baseline levels of urbanization, we stratified our analyses by global biogeographic subrealms (N=52; Fig. S1). Subrealms represent an intermediate hierarchical level within the One Earth^*82*^ (https://www.oneearth.org/bioregions/) bioregionalization framework, grouping the 185 terrestrial bioregions into broader units that reflect shared species pools and ecological contexts while maintaining meaningful regional structure. This scale represents a practical compromise between analyzing data at the finer bioregion level (which would result in many regions with insufficient observations for robust analysis) and broader classifications such as continents or the 14 biogeographic realms, which aggregate ecologically distinct regions and species pools. This regionalization has been widely used in macroecological and biogeographic research to contextualize species–environment relationships because subrealms capture meaningful gradients in biotic assemblages that are not accounted for by climatic classifications alone^*83,84*^.

This stratification allows species’ associations with urban environments to be interpreted relative to the environments available within the regions they occupy. This is important, as previous work has shown that species’ responses to urbanization are constrained by biogeographic context, because regional species pools reflect shared evolutionary, ecological, and historical filters^*23*^. Previous work has also shown that urban associations among species are context-dependent, and interpreting species’ responses without accounting for regional baselines conflates availability of urban environments with species’ affinity to them. This distinction is critical because identical levels of urbanization (e.g., VIIRS radiance) can have different ecological meanings across regions with different species pools and land-use histories. It avoids conflating species’ urban affinity with global differences in urban availability.

### Calculating urban affinity

After each GBIF occurrence record was assigned a VIIRS radiance value, as described above, we summarized these values at the species level within each subrealm. Specifically, for each species *s* within each subrealm *r*, we calculated the mean VIIRS radiance across all occurrence locations of that species in that subrealm. We refer to this quantity as the subrealm-specific urban score (𝑈_𝑠,𝑟_). Formally, the urban score for species *s* in subrealm *r* is defined as:

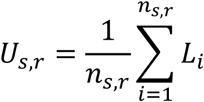

where 𝑛_𝑠,𝑟_ is the number of GBIF occurrence records for species *s* within subrealm *r*, and 𝐿_𝑖_ is the VIIRS night-time lights radiance value associated with occurrence *i*. The urban score is therefore an absolute descriptive summary of the urbanization levels associated with a species’ occurrence locations within a given subrealm. Higher urban scores indicate that a species tends to be observed in more highly urbanized (i.e., more brightly lit) environments, whereas lower values indicate occurrence in less urbanized environments. These urban scores serve as the intermediate step in our workflow and form the basis for the subsequent calculation of urban affinity.

To express species’ urban associations relative to the regional context in which they occur, we converted species-specific urban scores into a subrealm-specific measure of urban affinity. This step accounts for differences in baseline urbanization among regions and ensures that species are evaluated relative to the environments available within their biogeographic context. Within each subrealm r, we defined urban affinity for species s as: Aₛ,ᵣ = Uₛ,ᵣ − mean(Uᵣ), where 𝑈_𝑠,𝑟_ is the species-specific urban score calculated for species *s* within subrealm *r*, and 𝑚𝑒𝑎𝑛(𝑈_𝑟_)is the mean VIIRS radiance across all occurrence records of all species in that subrealm. This transformation centers species’ urban scores on the regional background level of urbanization.

Urban affinity values therefore describe whether a species tends to occur in environments that are more urbanized or less urbanized than is typical for that region (Fig. S6). Species with negative values occur disproportionately in less urbanized environments (urban avoiders), whereas species with positive values occur disproportionately in more urbanized environments (urban exploiters). By default, these values are relative measures, interpretable only within subrealms, and are not intended to represent absolute or globally comparable levels of urbanization. Importantly, this metric quantifies a species’ realized spatial association with urban environments relative to the regional background based on occurrence data. This framing is consistent with previous macroecological studies that infer species’ environmental affinities from spatial distributions rather than performance metrics (e.g.^*21,85*^). Consistent with this interpretation, previous work has also shown that measures of urban affinity (sometimes referred to as tolerance) calculated from VIIRS night-time lights are strongly correlated with analogous metrics using alternative proxies of urbanization such as human population density and the global human modification index^*45*^.

We applied the above procedure to any species that had at least 100 observations in a subrealm. Previous work has shown that this cutoff approximates the variability of a species’ response to urbanization and ensures that enough of a species urban habitat use has been captured^*21,86*^. After this filtering we were left with a total of 56,181 data points—i.e., unique urban affinity measures of a species in a subrealm—for potential inclusion in our analyses. These records represented 30,373 species that had at least one urban affinity measure, from 47 classes and across 51 subrealms (Fig. S7; Data S1). The number of subrealms a species was found in was mostly 1 (64%) but ranged from 1 to 34 (Fig. S8). All urban affinity measures (i.e., urban affinity by subrealm measures) which we produced as part of our workflow are made available, as we hope to advance further testing of urban affinity among different taxonomic groups and regions, while highlighting that correct interpretation of these relative urban affinity values is essential before use.

Sampling biases are common in large biodiversity databases, including GBIF. For instance, urban areas are better sampled compared to remote regions, as contributors typically concentrate around areas with high human density. However, our approach mitigates this concern by calculating each species’ urban affinity relative to the regional background of observations within the same subrealm. As a result, any broad-scale bias toward sampling in human-dominated landscapes affects all species within a region similarly, allowing meaningful comparisons among species rather than reliance on absolute estimates of urban occurrence. Importantly, our interpretation therefore focuses on relative differences among species within shared geographic contexts, rather than on absolute levels of urbanization. Previous work has found that this approach of assigning urban affinity is strongly correlated with occupancy-detection models where the target-background sampling is explicitly accounted for (see Figure S5 in ^*21*^). As illustrative examples, these data can be used to better understand the differences in urban affinity among Hymenoptera in Northeast American Forests (Fig. S9), Lepidoptera in Southeast Asian Forests (Fig. S10), or Asterales in Scandinavia & West Boreal Forests (Fig. S11).

### Species urbanness distributions

Our analysis of species urbanness distributions was descriptive in nature, relying on data visualization using ggplot2 to make histograms and density histograms of urban affinity values for each subrealm. In these analyses at the subrealm level, species are treated equally without any differentiation taxonomically, compared with the taxonomically-stratified analyses where species within each taxonomic rank are treated as similar (see below). To evaluate the consistency of SUD distributions across taxonomic levels and regions, we compared distributions among the most data-rich subrealms using violin plots for groups represented by ≥50 species (Supplementary Figures S2–S4).

### Aggregating measures of body size

Our objective was to compile a dataset of species-specific measures of ‘body size’, or ‘organism size’ more broadly to integrate with our measures of species-specific urban affinity described above. We aimed to find measures for as many species as possible, maximizing our sample size for modeling. For plants, we used ‘plant height’ as a proxy for body size, and data were downloaded from the TRY database—a database compilation of plant traits from many different dataset^*87*^. For animals, body size can be measured in different ways, depending on the taxonomic group of interest (e.g., body length, wingspan, biomass, shell size, radius, forewing length, body mass, or biovolume). Therefore, our dataset was purposefully broad in its composition of body size estimates. Because ‘body size’ is the predominant form referred to in the literature, we use this throughout, but take it to broadly mean ‘organism size’ as discussed above.

To build the animal dataset, we first performed a semi-systematic broad literature search, using Web of Science, Google Scholar, Zenodo, Dryad, and Figshare with variations of the key words: “body size”, “organism size”, “body length”, “body mass” and “interspecific”, “global patterns”, “animals”, and higher-level groups such as classes (e.g., “Aves”, “Birds”, “Mammals”, “Mammalia”). We started by downloading known compilations for various taxa, for example mammals^*88*^ or Odonata^*89*^. In these instances, we spent minimal further time searching for specific body size data for these larger taxonomic groups, assuming that these compilations had already aggregated the majority of data available. Following this, we conducted more detailed literature searches, predominantly using Google Scholar and Web of Science, which focused on our *a priori* list of species for which we had a measure of urban affinity (Data S1). These were conducted at class, order, or family taxonomic levels, but not at taxonomic levels below family. Multiple keywords were used for each group while searching when it was obvious to do so. For example, when searching for body size measures in Coccinellidae both “ladybirds”, “ladybugs”, and “Coccinellidae” were used in search strings, as well as “wing length”, “body length”, or “body size”. Search effort was qualitatively proportional to the number of species we had in each taxonomic level. For example, because we only had 43 species with urban scores from the class Diplopoda but 3,515 species from the order Lepidoptera, we spent proportionately more time and effort searching for body size data for Lepidoptera. Similarly, we spent more time on hard-to-find taxa, namely groups from Insecta. In our searches we generally ignored studies that were focused on intraspecific variation in body size for one or a few species, unless we found that the species was included in our urban affinity list. In addition to literature searches, we emailed networks of colleagues, looked at reference lists in papers that had data, and emailed corresponding authors of papers who had potentially relevant data but for which data were not accessible.

After this semi-systematic searching, we re-assessed the remaining species for which we had an urban affinity measure but not a measure of body size. We then performed a second-level targeted search (i.e., more systematic) for any family with at least 10 species remaining for which we had a measure of urban affinity. First, we searched Web of Science and Google Scholar one family at a time and looked at the first 100 hits, with the family name ‘and’ “body size” used as search terms. Second, we used Google search engine to search for these remaining species individually (N∼3,100). We focused on including species for which Google immediately returned an estimate of body size (e.g., through Wikipedia, or an aggregating website), and did not go to original species descriptions. This detailed searching and the aggregation of body size measures concluded in June 2022.

### Measurements of body size

In our searching for body size, we were agnostic to the type of body size measure, given that many different measures are used for quantifying body size, or proxies for body size. For each dataset we downloaded or processed, we kept track of the following key metadata properties: ‘type’, ‘units’, and ‘measurement detail’. Type refers to the different types of body size measurement, ranging from body length and body mass to Weber’s length in ants. Table S1 shows a list of the different body size types encompassed in our analysis (see statistical details below for how different body size types were accounted for in our analyses). For each dataset, we also kept track of the units (e.g., mm, g, mg cm^-2, or mm^3). And lastly, wherever possible, we noted the measurement detail of the body size measures, whether the mean, median, or maximum was used, or whether it was for males or females only. Many times, however, this information was not readily available, and thus this metadata field was not required in our dataset composition.

### Taxonomic harmonization and dataset integration

Because body size data were aggregated across many potentially different datasets, we performed a taxonomic harmonization step to regain some species that did not match due to typos, spelling mistakes, or synonyms. For this, we treated Plantae and Animalia separately and used the taxize package^*90*^ in R to search through the list of species and find any synonyms or fuzzy matches of other taxonomic entities, always matching back to the GBIF taxonomic backbone, given that our urban affinity measures follow this taxonomy (Data S1). With the possible synonyms and matches, we then integrated the urban affinity measures with the body size dataset using either direct matches or matches of synonyms. The taxonomic harmonization step regained a total of 831 species (see Fig. S12) that were integrated in the dataset for analysis. Data S3 provides details on the 223 different ‘datasets’ (i.e., downloaded data, or manually added data) that were processed as measures of body size. The most prevalent class in our final dataset (Fig. S13) was Insecta (N=7,526), followed by Magnoliopsida (N=4,872) and Aves (N=3,313).

Our final dataset for potential analysis and modeling (Data S2) included a total of 94,087 observations (i.e. unique combination of species urban affinity values, subrealm, and body size measure) for 20,957 species that had at least one measure of urban affinity and at least one measure of body size.

### Quantifying the relationship between urban affinity and body size

Our objective here was to understand the relationship between urban affinity (the response variable) and body size (the predictor variable). Our dataset had potentially multiple measures of body size for an individual species, and differing levels of metadata. For example, some datasets may have specified whether the mean was used, or others might have specified for females only (see Table S1). Therefore, before modeling, our goal was to minimize the undue effect of having too many random effect levels. First, because an individual species could have multiple measures of body size, we selected the body size measure that was most common among the species in each family. For example, if a species had both a measure of body length and of body mass, we selected the measurement type that was most commonly available among the species in each family. Second, we further reduced the potential number of levels and variation in body size measurement by creating a synthetic “metadata” variable that was the combination of ‘type’, ‘units’, and ‘measurement detail’. By doing this, we decreased the potential number of levels of a random effect from 223 (i.e., the number of data sources) to 71 (i.e., the combination of type, unit, and measurement detail). Third, several papers/data sources could have all reported the same metadata (e.g., maximum body length for females in mm), and instead of being treated as individual random effects because they were from different data sources, we assumed that these represented similar data and collapsed them into one random effect level (here, for maximum body length for females in mm). Importantly, this procedure did not result in the exclusion of species lacking a particular body size measurement type; rather, all species with at least one available body size estimate were retained, with measurement heterogeneity explicitly accounted for through hierarchical modeling.

### Overall kingdom-level model construction

We used Bayesian hierarchical models to test for the effect of body size on urban affinity. In our models, urban affinity was the response variable and body size was the predictor variable of interest. To evaluate the overall relationship between body size and urban affinity, we fitted two comprehensive models that incorporated hierarchical taxonomic levels and the relationship with body size. The first model included fixed effects for kingdom and log-transformed body size due to its skewed distribution, along with their interaction, to investigate differences in the body size effect on urban affinity between plants and animals. The random effects accounted for hierarchical taxonomic levels, incorporating random intercepts and slopes for class, order, and family to capture variation at each level. Additional random effects for metadata and subrealm were included as random intercepts to control for study-specific differences in data and regional differences in urban affinity. The model formula was:

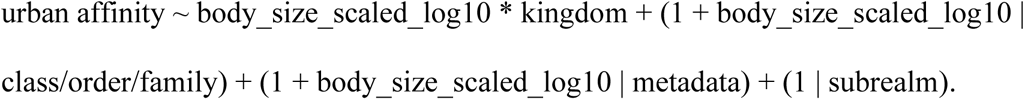

The second model built upon this structure but only included an interaction term between log-transformed body size and kingdom as well as kingdom as a fixed effect to determine if the effect of body size on urban affinity varied consistently across the plant and animal kingdom. It maintained the same hierarchical random effects structure as the first model, with class, order, and family nested to account for hierarchical variation, and metadata and subrealm included as additional grouping factors. The formula was:

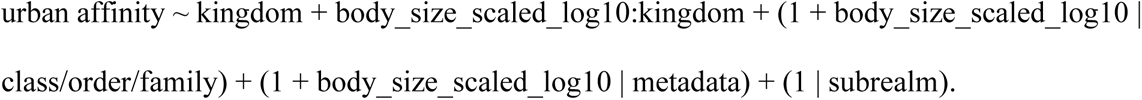

### Family-specific model construction

For our main analyses we chose to employ a ‘many models’ approach, where we fit individual models for each family that met a set of minimum criteria because many families do not have a known association or published information on the extent to which urban affinity associates with body size. A family was only included if there were at least 10 unique species with an urban affinity measure and body size measure, and a subrealm was only included if it had at least two species included to specify a random slope. For example, if there was only one species of a particular family included in our dataset, this family would not have received an independent model. We decided to focus our main analysis at the family level for three key reasons. First, for most of the taxonomic families included in our analyses there is little *a priori* expectation about the effect of body size on urban affinity (i.e., little to no literature on the subject; but see McClain and Boyer^*39*^), and therefore we wanted each model to represent an advance in knowledge of that relationship for that family. Second, there were large imbalances in the number of data points available for each taxonomic group, with some families having hundreds of species with available data and others with only a handful of species. Thus, we wanted to avoid shrinkage where the model results were primarily driven by the taxonomic groups with a larger number of data points represented. Third, we wanted to avoid an instance where a family (or other taxonomic grouping) could have little or no relationship between body size and urban affinity, but if included together in one model could result in a positive effect (see Fig. S14 for a simulated example). This was a smaller concern at the family level, which we present as the main analysis, because of the smaller imbalance in data availability across the taxa within each family.

When we fit the family-specific models, there were different requirements for the random effect structure depending on how many subrealms and metadata sources (i.e., types of body size measures) were available. In the simplest case, all species in a family were just from one subrealm and had just one data source for body size. But this was not always the case, and therefore we developed four different ‘types of models’ which varied in their treatment of random effect structure. Each ‘type’ of model, however, had the same overall objective—to quantify the relationship between body size and urban affinity. The four model types are as follows. Type 1: had at least two subrealms and at least two types of synthetic metadata (see above) sources included. Type 2: had data from only one subrealm and only one type of synthetic metadata source included (but could have data from more than one initial data source). Type 3: had at least two subrealms and only one type of synthetic metadata source included. Type 4: had data from only one subrealm and at least two types of synthetic metadata sources included. For all types of model fits, the response variable was the urban affinity measure, described above, and the fixed effect of interest was body size which was log10-transformed and scaled and centered by the synthetic metadata source. Type 1 had a random intercept and slope for subrealm, allowing the intercept and slope to vary across different levels of subrealm as well as a random intercept for the synthetic metadata source. Type 2 had no random effects. Type 3 had only a random intercept and slope for subrealm, allowing the intercept and slope to vary across different levels of subrealm. Type 4 had only a random intercept for the synthetic metadata source. Modeling was done using the ‘brm()’ function in R from the brms package^*91,92,93*^. For all model types we specified a standard normal prior, with a mean of 0 and a standard deviation of 1 applied to the fixed effects of the model. Models were fit with 1000 warmup iterations, 6000 iterations for the Markov Chain Monte Carlo sampling, 4 chains, and the adaptation target acceptance probability was set to 0.99.

For models of type 1 and type 4, where we combined metadata for body size measurements among different species, we also explored more complex model structures that included a random slope for body size by metadata source, in addition to subrealm. Such a model structure would allow the relationship between body size and urban affinity to vary across measurement types. However, these models often substantially reduced interpretability and inflated uncertainty around slope estimates, especially when metadata groups were sparsely represented, which was common. Additionally, many of the families only had 2 types of metadata, limiting our statistical power to fit random slopes. Comparative analyses showed that such models produced weaker or more uncertain estimates than our main model structure (i.e., random intercept for metadata of body size). We also note that to account for heterogeneity in body size measurement types, all body size values were log10-transformed, scaled, and centered by their metadata source prior to analysis, following standard practices in allometric studies. We use two illustrative examples (Figures S15 and S16) to illustrate the influence of including a random slope for metadata source.

### Additional analyses beyond family level

Although our main analysis was focused on the family level, one potential drawback is that species could be excluded from any of the analyses for a family that has a small number of species. Plus, because some analyses have previously been conducted at higher taxonomic levels (e.g., Lepidoptera), we repeated the above procedure described for family at the order (see Supplementary Text) and kingdom level.

## Acknowledgements

We thank Tim Robertson and the GBIF support team in supplying avro files of the GBIF download (https://doi.org/10.15468/dl.9ufars) for upload to our personal Google Big Query. Henrique M. Pereira provided helpful discussion in the early stages of this project. We thank the following people for help in qualitatively assessing our proxy for urban affinity: Luis Aguirre, Timothy Bonebrake, Jason Bried, Alex Córdoba-Aguilar, Michael Crossley, Wouter Deconinck, Geert De Knijf, Tim Gardiner, Michael Greenfield, Ralf Gyselings, Frederik Hendrickx, Axel Hochkirch, Roy Kleukers, Anton Kristin, Paulo Lemos, André Victor Lucci Freitas, Onildo João Marini Filho, Daniele Matenaar, Elisa Montes-Fuentemayor, Tara Murray, Howon Rhee, Leslie Ries, Szabolcs Sáfián, Ann Swengel, Scott Swengel, Pieter Vantieghem, Wade Worthen. We thank the following people for supplying unpublished data of body sizes: Roel van Klink, Michelle Tseng, Erik Öckinger, Philip Barton, Werner Ulrich, Axel Höchkirch, Thomas Sherratt, Michiel Wallis de Vries, Lea Heidrich, Jörg Müller, Robert Tropek, Maldwyn Evans, Martin Gossner, Bernhard Hausdorf, Stano Pekar, Carlo Seifert, Onildo João Marini Filho, Sei-Woong Choi, Dominik Rabl, Markus Franzén, Elena Piano, Derek Hennen. CTC, DEB, IS acknowledge funding of iDiv via the German Research Foundation grant DFG FZT 118. CTC was supported in part by a Marie Skłodowska-Curie Individual Fellowship grant 891052 and was supported in part by the intramural research program of the U.S. Department of Agriculture, National Institute of Food and Agriculture, Hatch FLA-FTL-006297. VS was supported by Georgetown University, a U.S. National Science Foundation Graduate Research Fellowship grant 1937959, and a David H. Smith Conservation Research Fellowship. LHA was supported by the Academy of Finland grant 340280.

## Statement of authorship

CTC, DEB, TM conceptualized the project and developed project methodology. CTC, TM, VS, JHW, BMM conducted analyses. CTC conducted project administration and supervision. CTC, DEB, TM wrote the original draft, and CTC, DEB, VS, BMM, LA, IS, JHW, TM were involved in review and editing.

## Data accessibility

All data analysis was conducted in R statistical software and relied heavily on tidyverse (Wickham et al. 2019). The DOI representing our download is: https://doi.org/10.15468/dl.4dcbgt. Our processing of the GBIF data to quantify urban scores is available here: https://github.com/coreytcallaghan/body_size_vs_urban_tolerance. Not all body size data can be openly shared due to copyright restrictions, however the freely available dataset, cleaned and processed for our analysis is available here: https://github.com/coreytcallaghan/body_size_vs_urban_tolerance. In this repository we did provide reproducible data and code, albeit not for the entire dataset of body sizes. We will archive a cleaned version of this repository in Zenodo upon acceptance of this manuscript.

## Supplementary Materials

### Supplementary Text

In addition to family-level effect sizes, we performed the analysis at the order level, and found similar heterogeneous results. Among the 128 orders, a total of 45 had some evidence of an effect of body size on urban affinity (14 weak, 6 moderate, and 25 strong). Some of the orders with the strongest positive effect size included Sapindales (dicots, soapberry plants) and Columbiformes (birds, doves and pigeons), and orders with the strongest negative effect sizes included Coleoptera (insects, beetles) and Piciformes (birds, woodpeckers). Of note are the differential responses down the taxonomic guilds, where some families can show contrasting results, but there may be a strong effect when looking at the order level, or higher (e.g., Fig. S2).

**Fig. S1:**
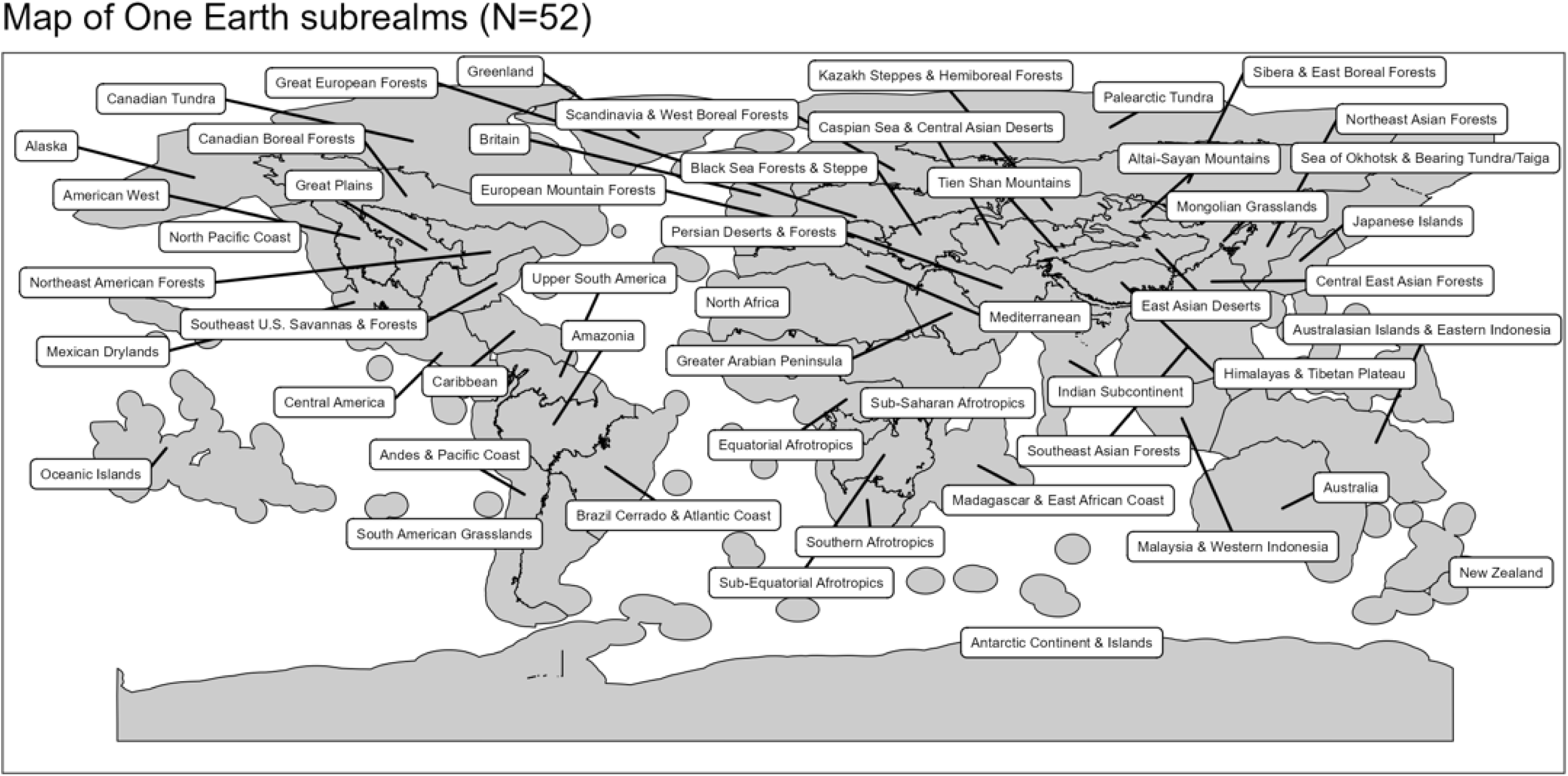
We used subrealms as our geographical aggregation. Subrealms were quantified from aggregating bioregions as identified by One Earth (see more here: https://www.oneearth.org/bioregions/). The subrealms level was chosen after exploring the tradeoff between accounting for geographic differences in urban affinity and the number of species that could be included.

**Fig. S2.**
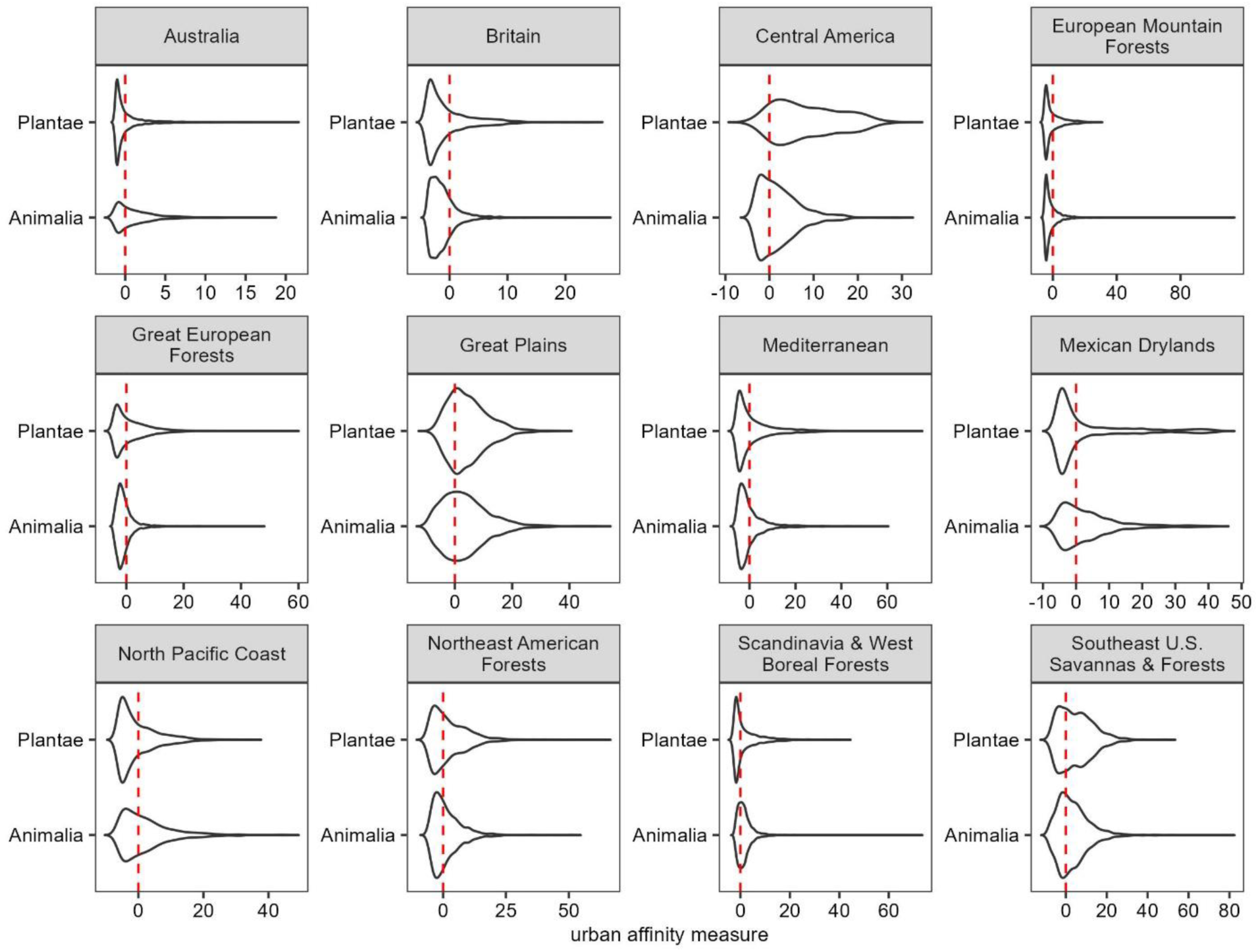
Violin plots showing the distribution of urban affinity values by kingdom across the 12 subrealms with the greatest sample sizes.

**Fig. S3.**
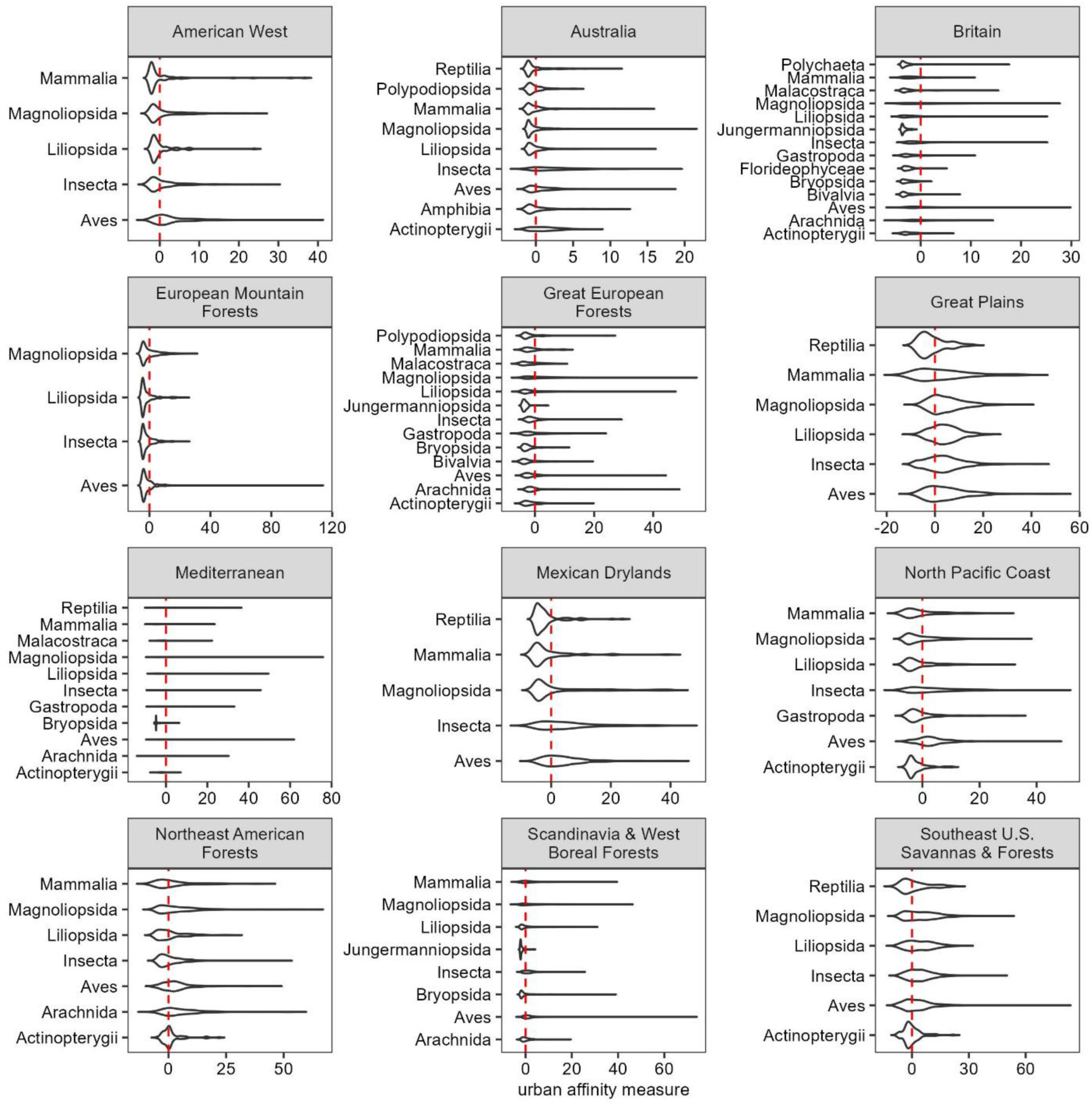
Violin plots showing the distribution of urban affinity values by class across the 12 subrealms with the greatest sample size. Only classes represented by ≥ 50 species within a subrealm are included.

**Fig. S4.**
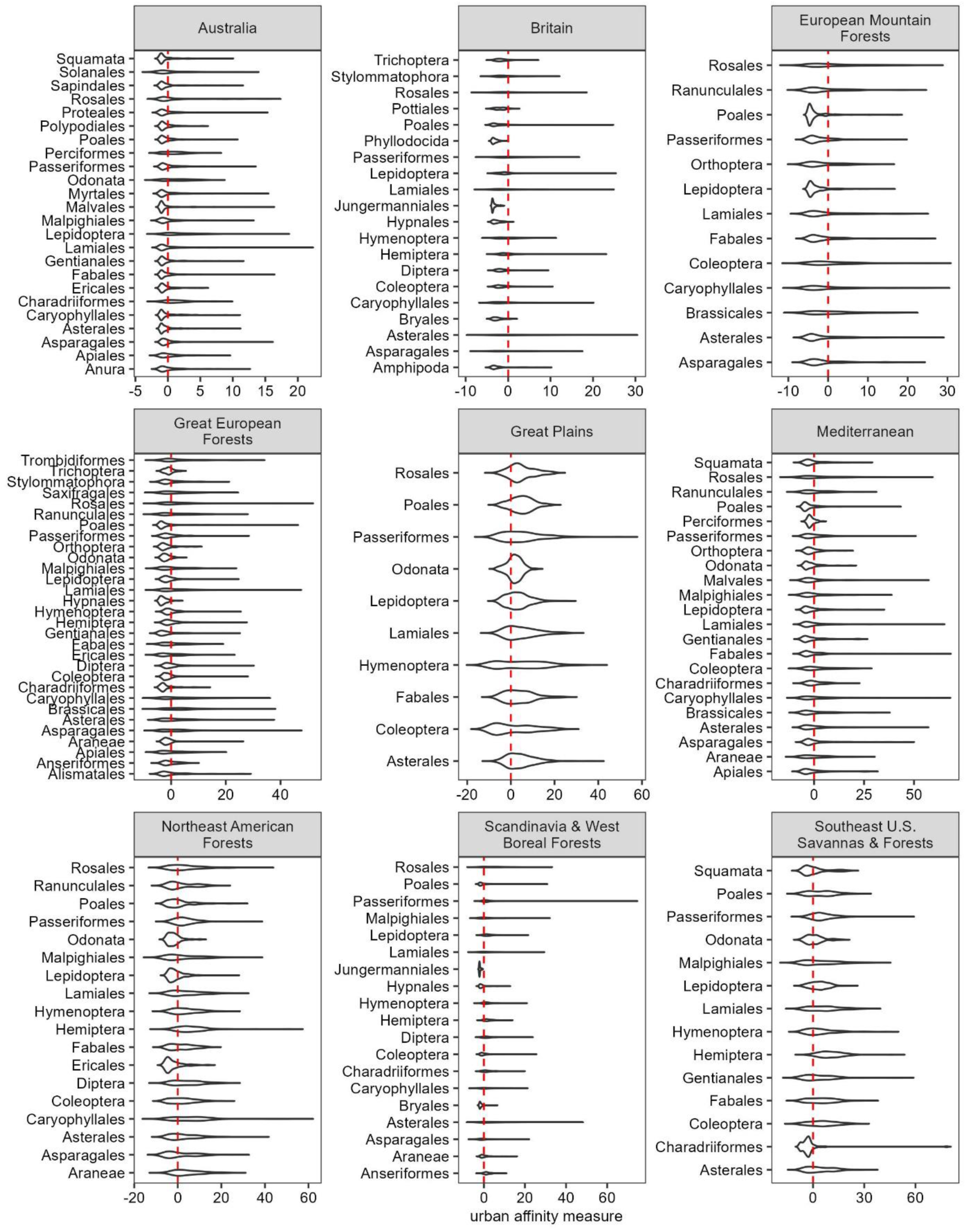
Violin plots showing the distribution of urban affinity values by order across the 9 subrealms with the greatest sample size. Only orders represented by ≥ 50 species within a subrealm are included.

**Fig. S5:**
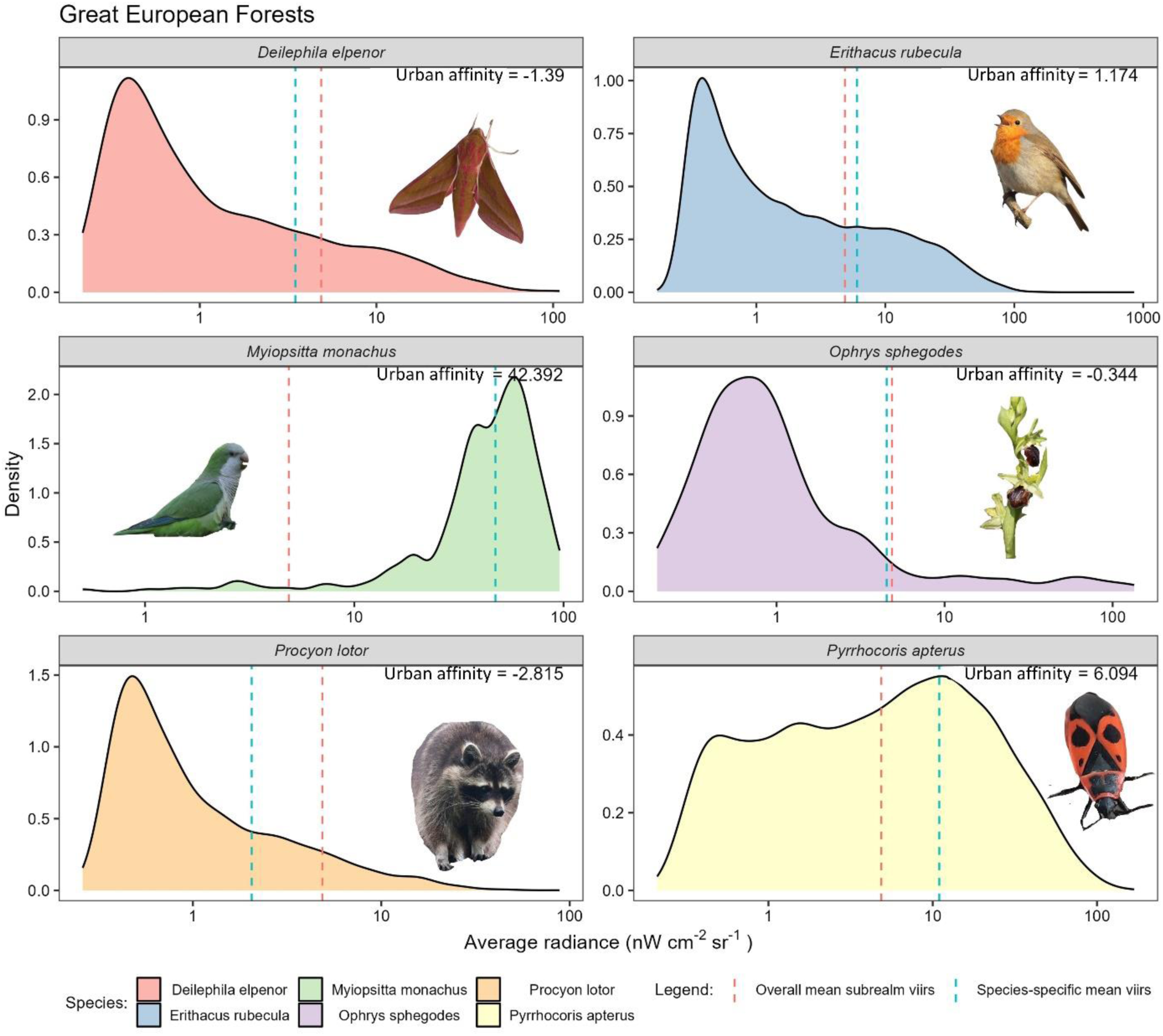
An illustration of the species-specific distribution of observations and the VIIRS values, in average radiance, of those observations for six different species within the Great European Forests subrealm. Of note is that the urban affinity is not shown, as some are negative values, but these values are shown in the text.

**Fig. S6:**
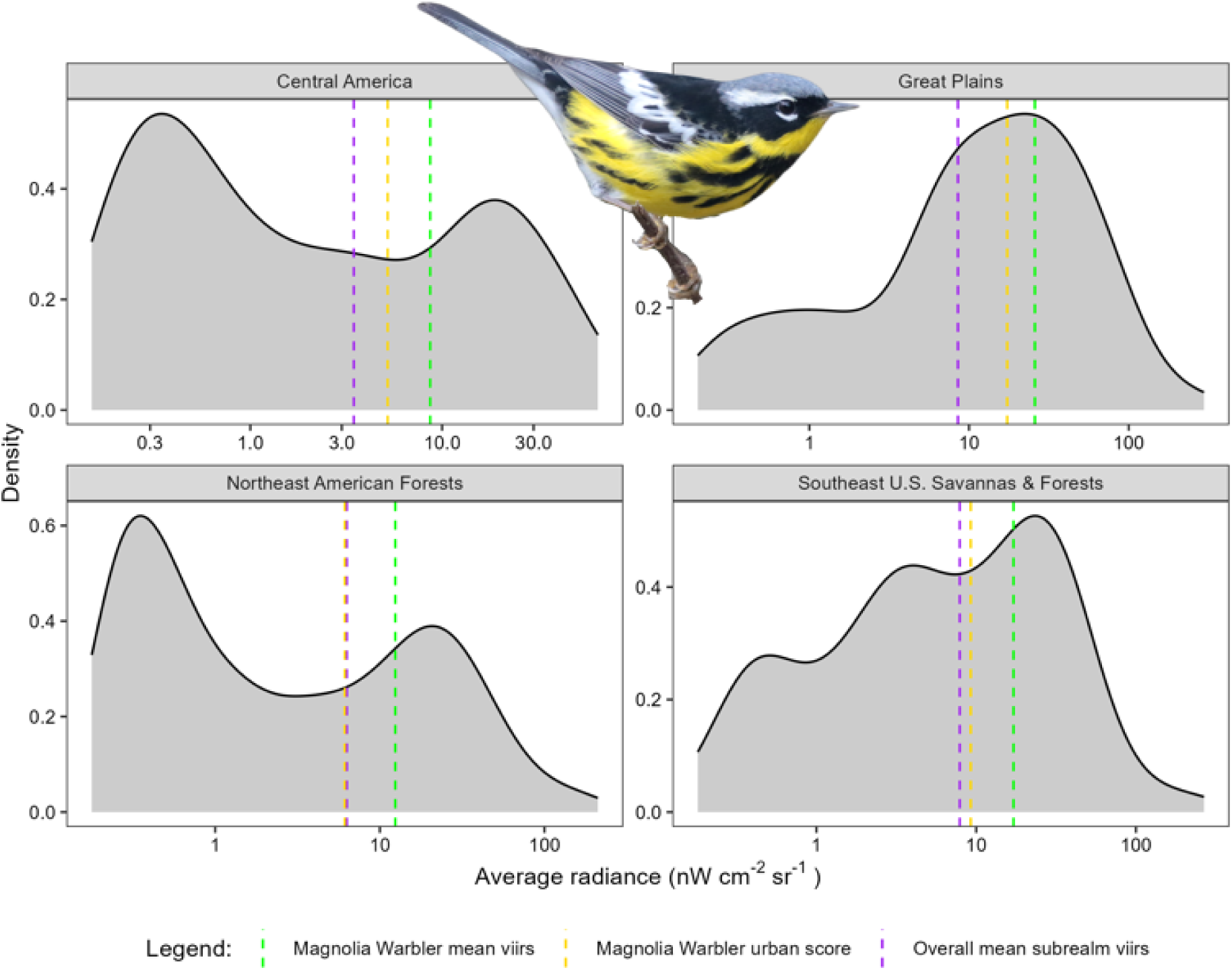
The distribution of Magnolia warbler *Setophaga magnolia* observations and the VIIRS values, in average radiance, of those observations in four different subrealms. Shown using dashed lines are the species-specific mean (green), the subrealm-specific mean of VIIRS (violet), and the resulting urban affinity measure (yellow).

**Fig. S7:**
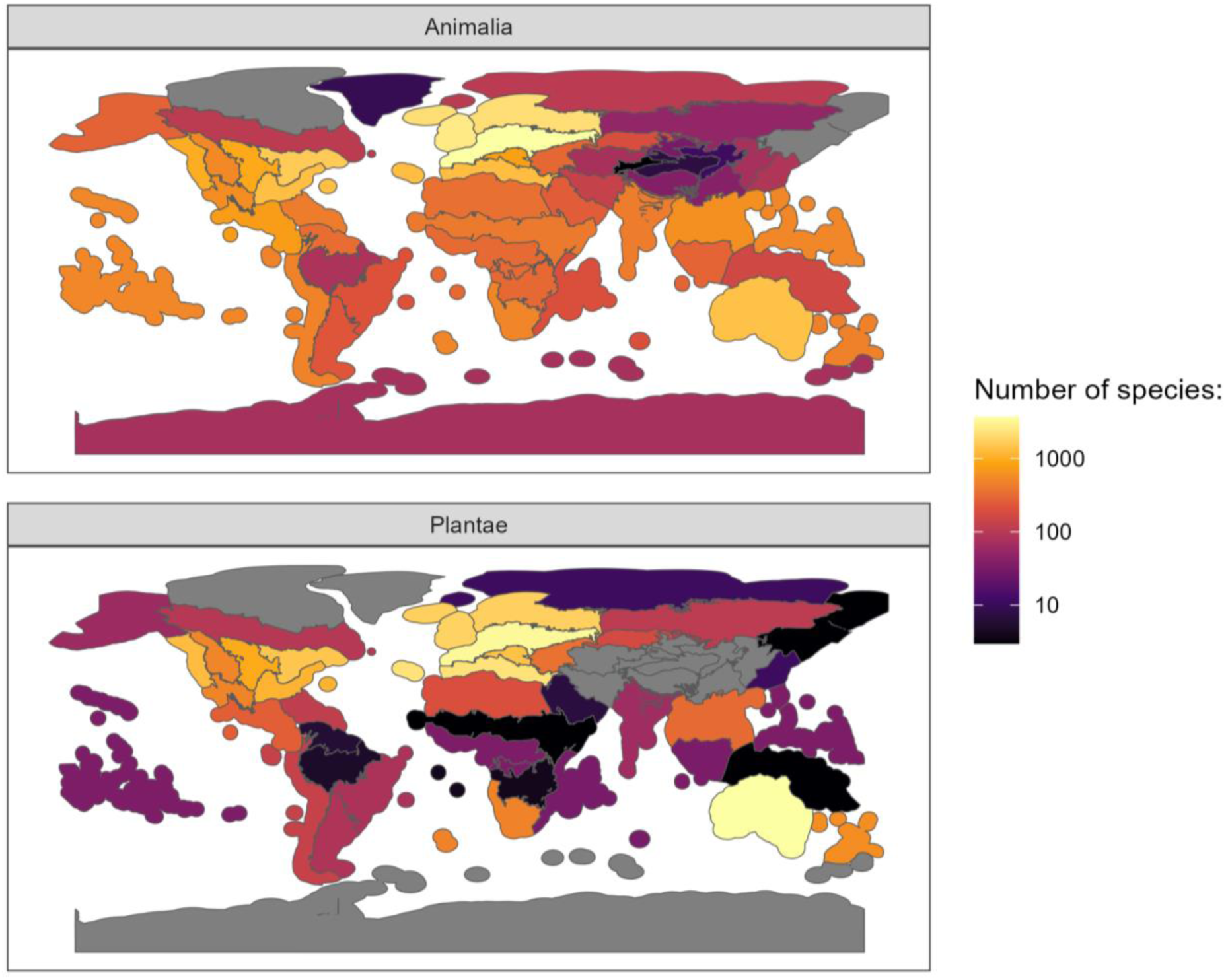
The total number of species for which we calculated urban affinity scores, stratified by subrealm and shown separately for Animalia (top) and Plantae (bottom). The subrealms with their names are shown in Figure S1. The dataset of species’ urban affinity scores is provided in Table S1.

**Fig. S8:**
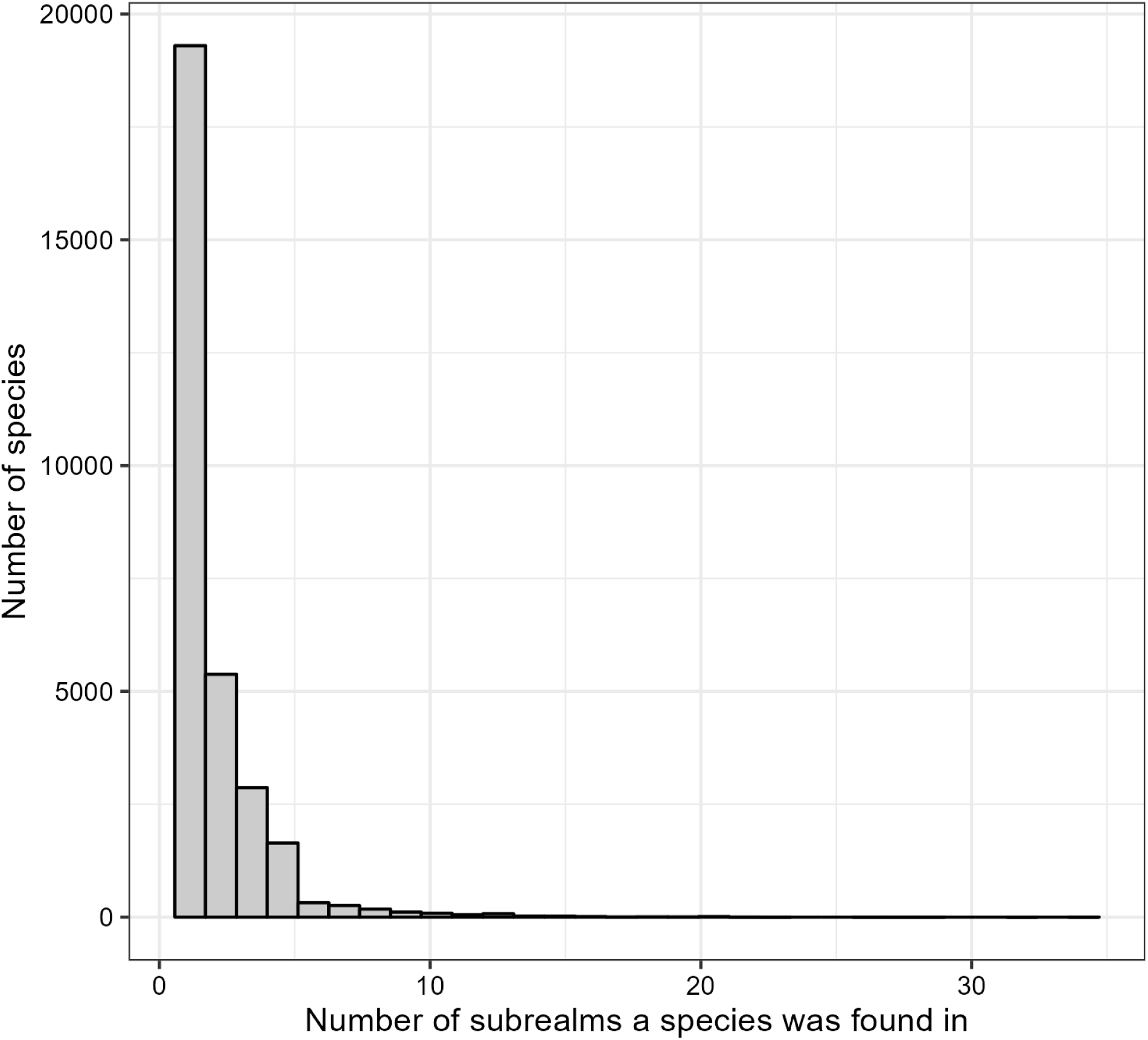
The total number of subrealms for which a species had an urban affinity score. The majority of species (64%) only had an urban affinity score from one subrealm, but the range was from 1 to 34.

**Fig. S9:**
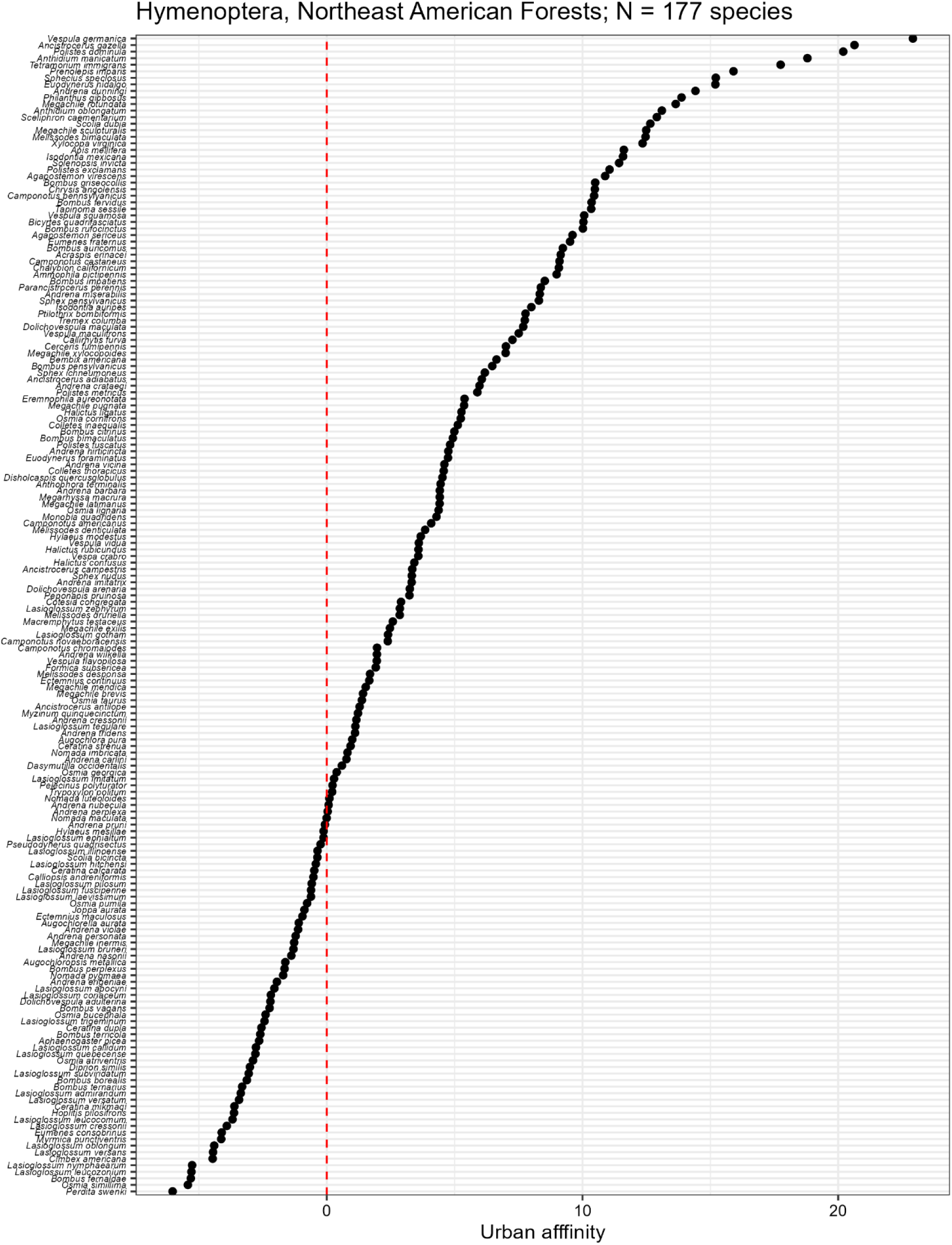
An illustrative example, showing the relative urban affinity scores for Hymenoptera in the Northeast American Forests subrealm. The plot is for illustrative purposes and the values for each species can be found in Data S1.

**Fig. S10:**
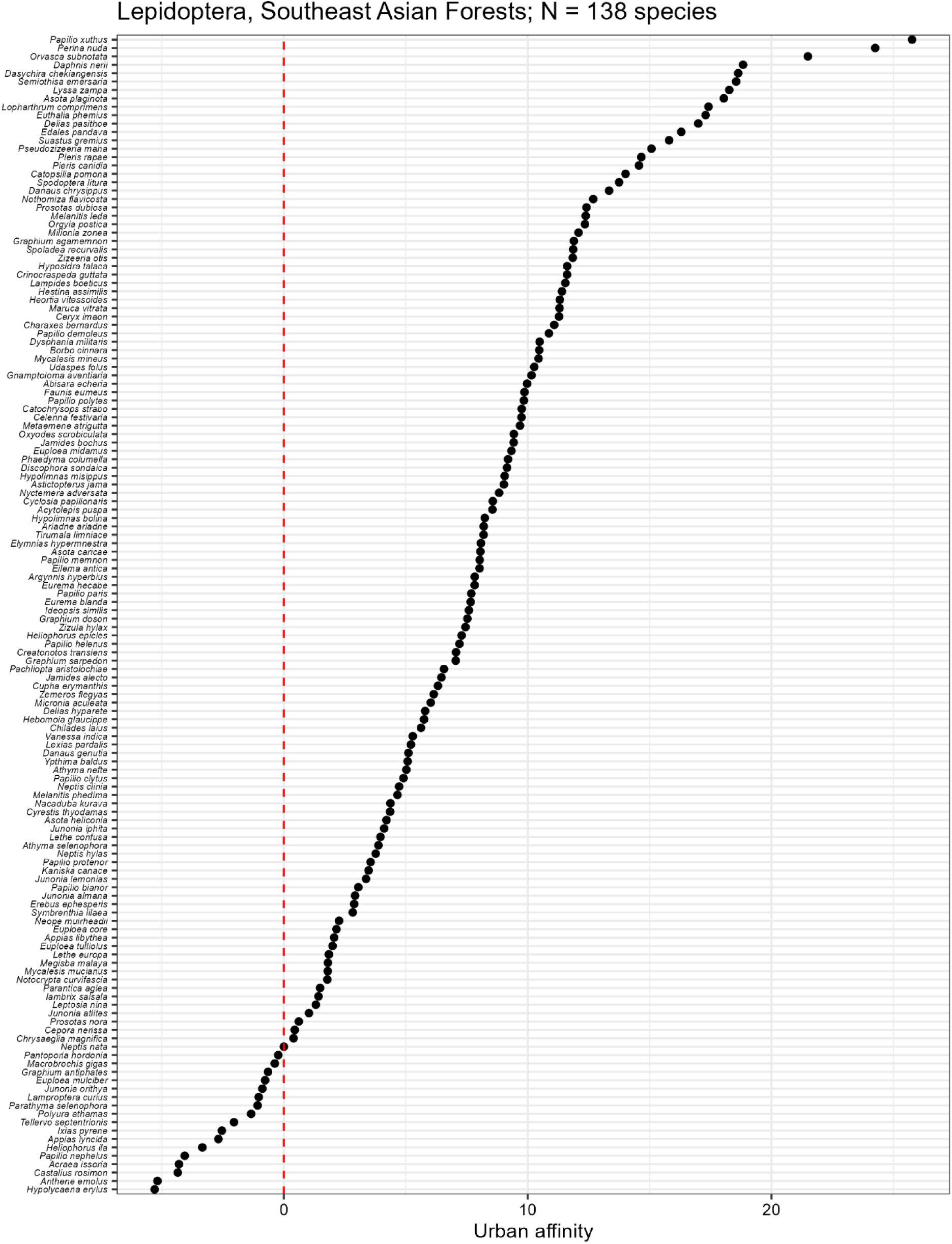
An illustrative example, showing the relative urban affinity scores for Lepidoptera in the Southeast Asian Forests subrealm. The plot is for illustrative purposes and the values for each species can be found in Data S1.

**Fig. S11:**
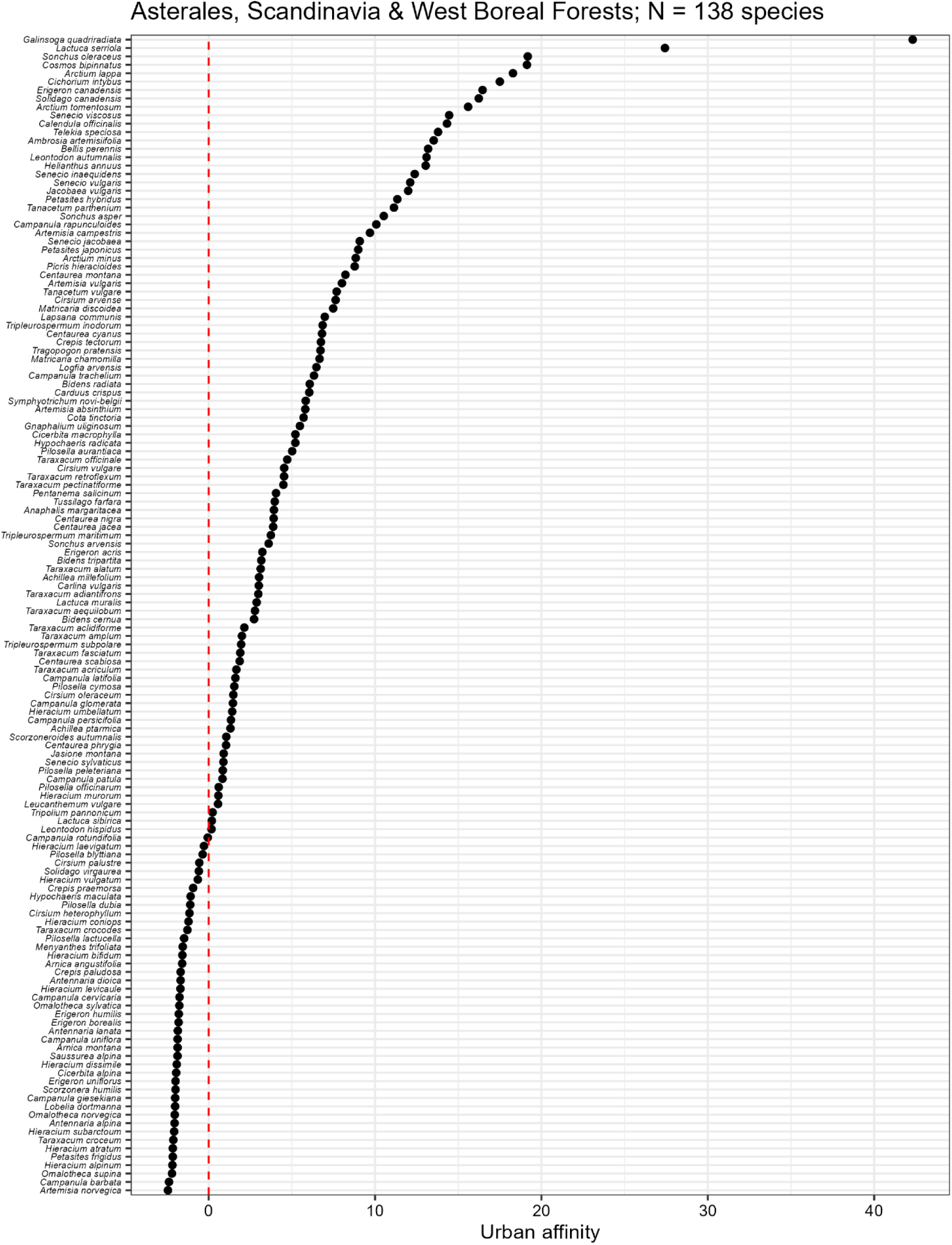
An illustrative example, showing the relative urban affinity scores for Asterales in the Scandinavia & West Boreal Forests subrealm. The plot is for illustrative purposes and the values for each species can be found in Data S1.

**Fig. S12:**
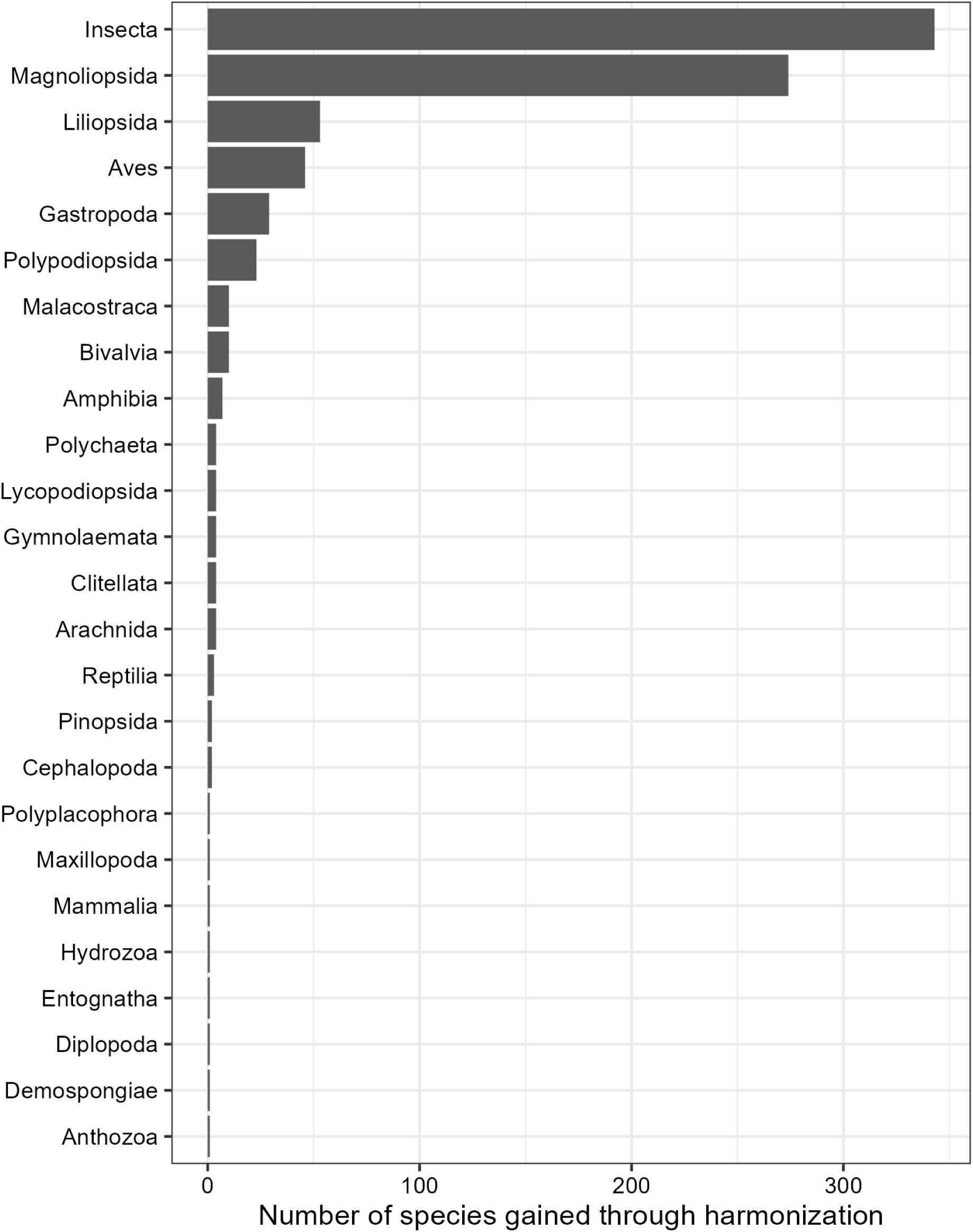
A summary of the number of species, shown per class, gained through taxonomic harmonization, and therefore included in the analysis dataset.

**Fig. S13:**
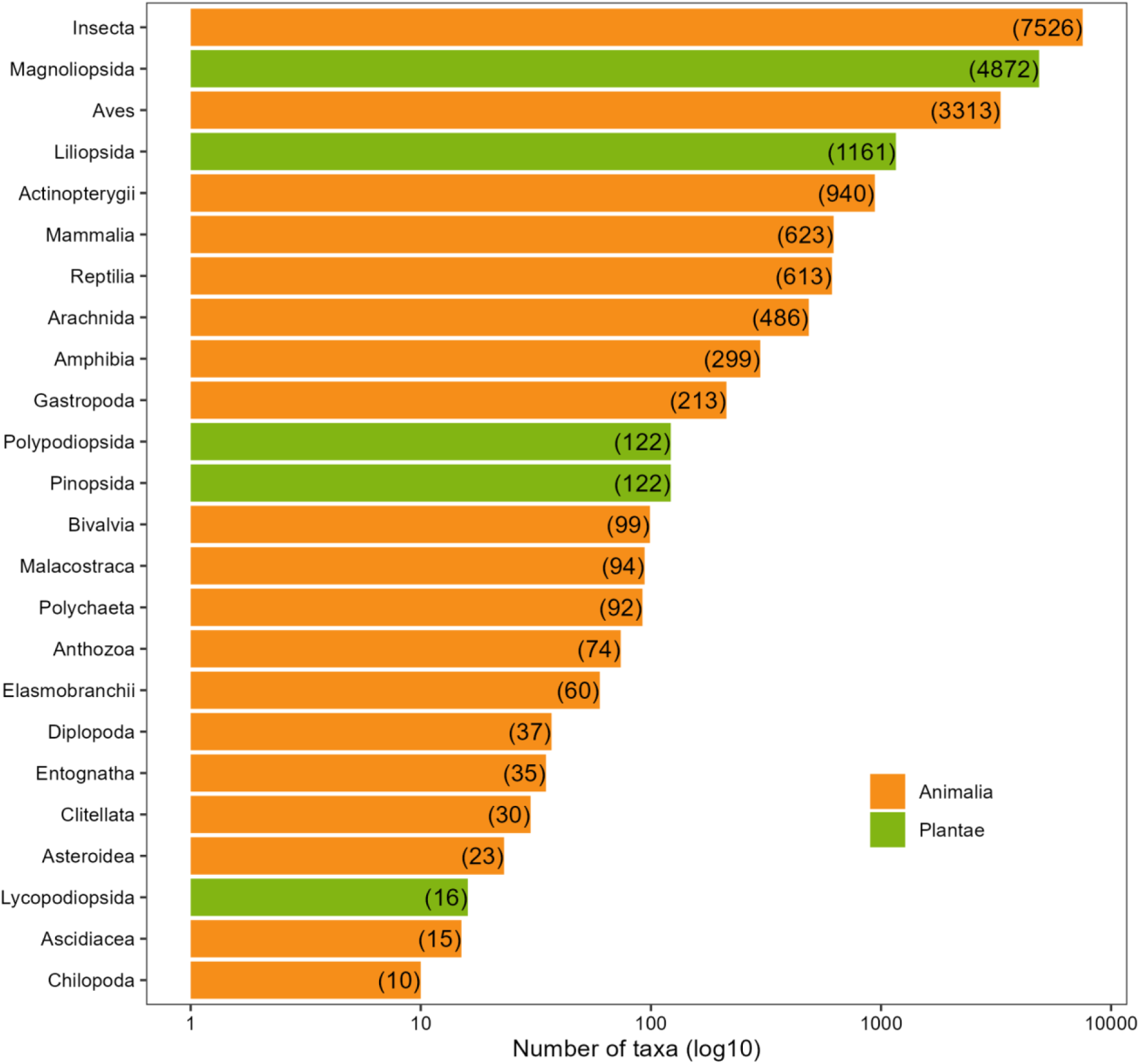
The number of taxa, on a log10-transformed scale. These can be found in Table S3.

**Fig. S14:**
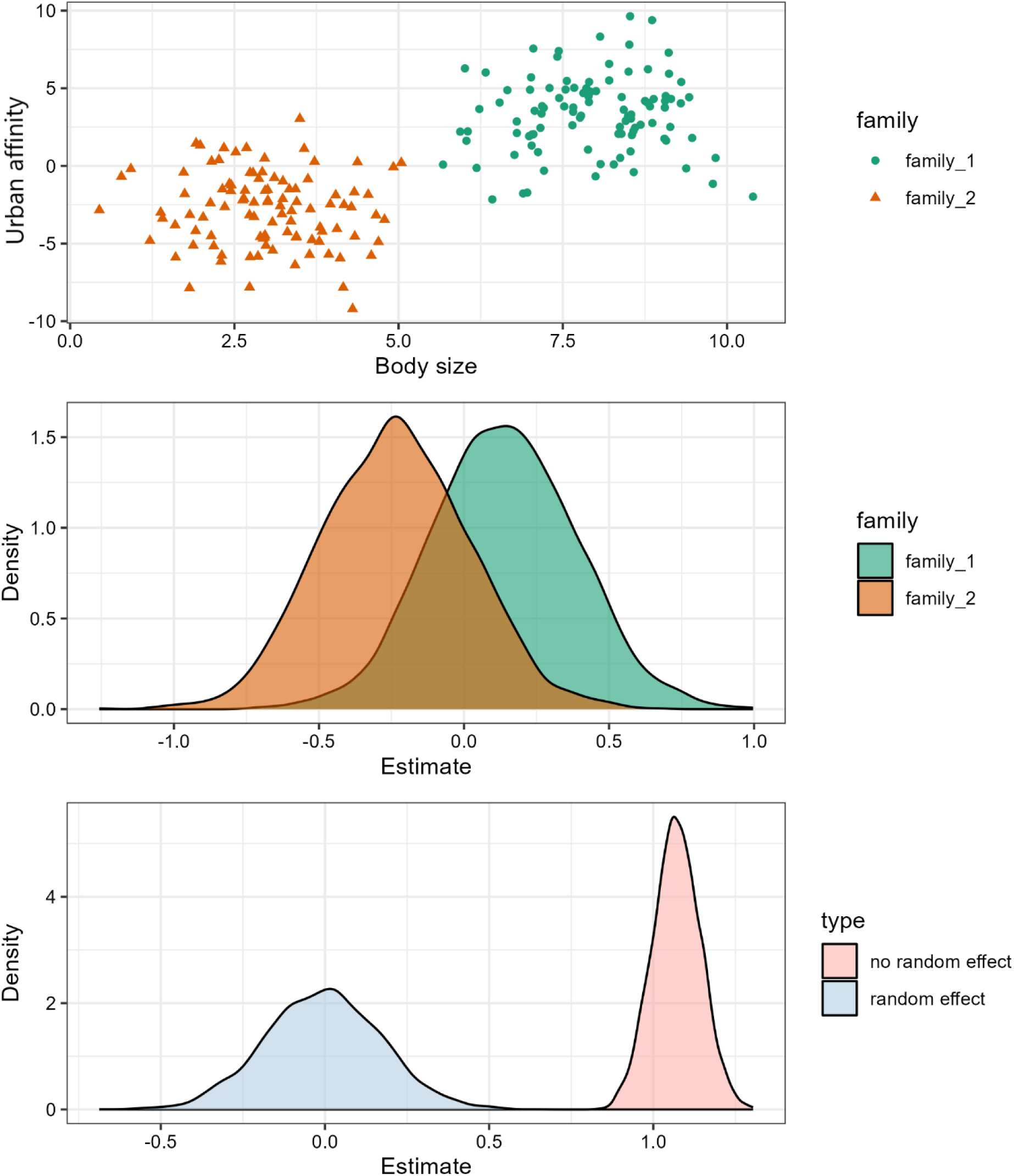
We chose to model each taxonomic group (i.e., family, order, class, phylum, and kingdom) independently of one another to avoid the influence of where individual families could have no effect but result in a positive effect if modeled jointly. The top panel shows simulated raw data for two families; the middle panel shows the posterior distribution of a bayesian model fit for each of these families separately; the bottom panel shows the posterior distribution of a bayesian model fit for a model including family as a random effect and for a model not including family as a random effect.

**Fig. S15:**
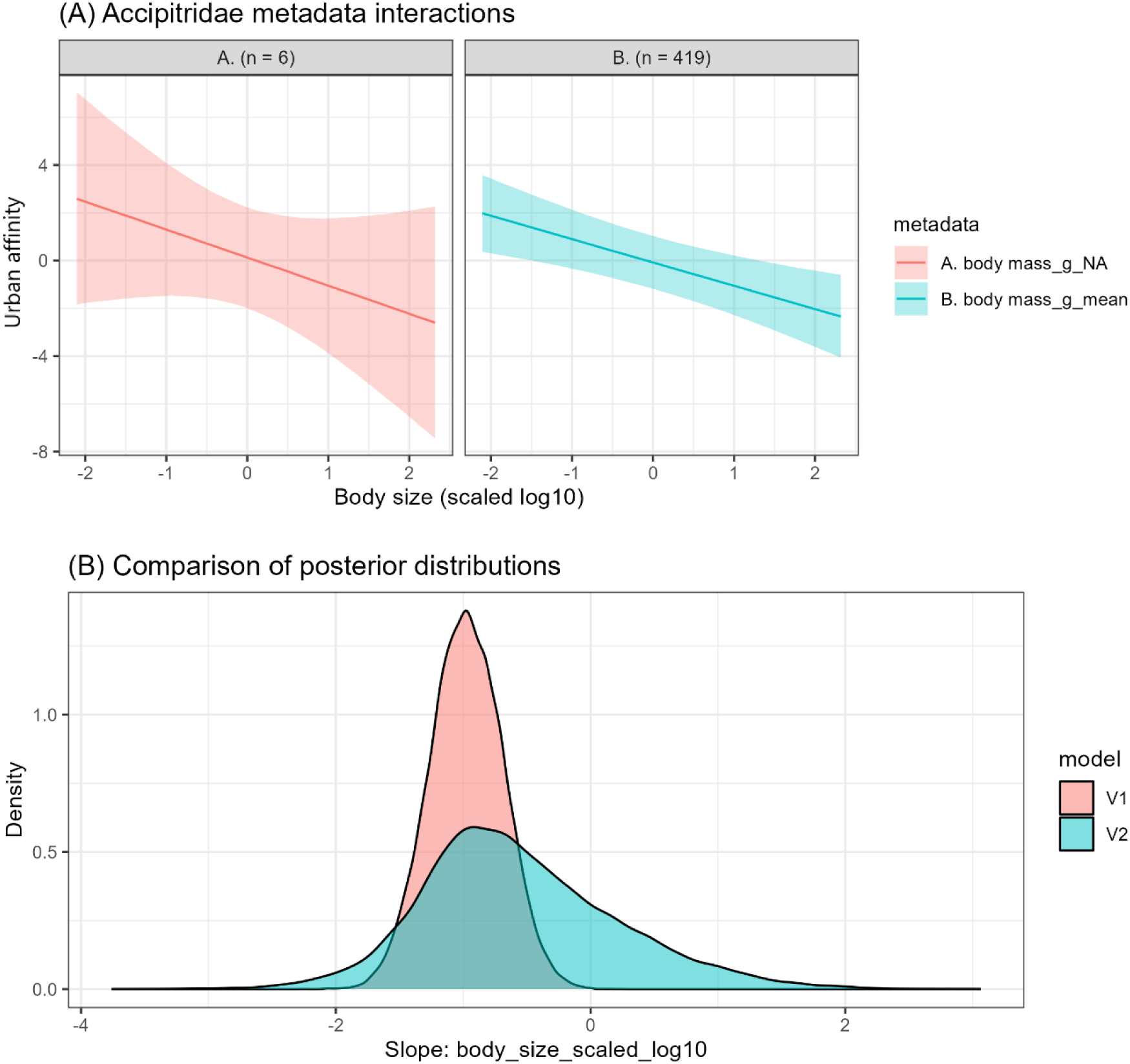
An illustrative example showing the influence of using a random slope for body size, by showing the (A) Interaction between body size (scaled and log10 transformed) and body size measurement type (metadata; see Methods for details) for the family Accipitridae. Facets represent different types of body size metrics used in the dataset, labeled with a letter identifier and corresponding sample size (n). Lines represent predicted urban affinity based on body size, with 95% credible intervals shown as shaded ribbons. These are extracted from a model fit with an interaction between metadata and body size to purposefully investigate the influence of metadata and the relationship with body size. (B) Posterior distributions of the slope for body size (scaled and log10 transformed) under two model structures: V1 includes a random slope for body size by subrealm and a random intercept for metadata (as presented in main results), while V2 adds a random slope for body size by metadata. The inclusion of this additional random slope in V2 increases uncertainty and pulls slope estimates toward zero, particularly when data are sparse within body size measurement types as illustrated here. Also see Fig. S15.

**Fig. S16:**
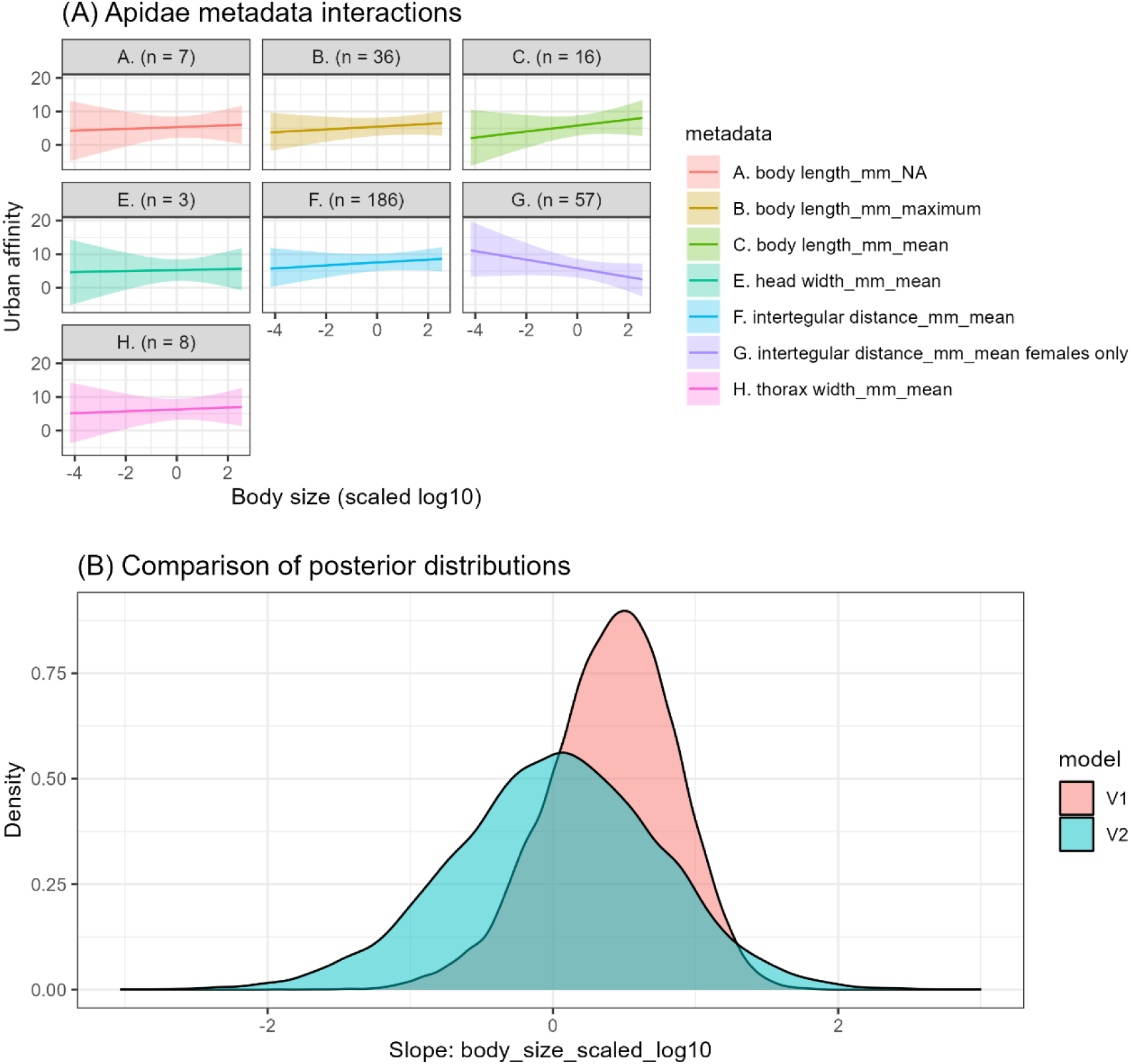
An illustrative example showing the influence of using a random slope for body size, by showing the (A) Interaction between body size (scaled and log10 transformed) and body size measurement type (metadata; see Methods for details) for the family Apidae. Facets represent different types of body size metrics used in the dataset, labeled with a letter identifier and corresponding sample size (n). Lines represent predicted urban affinity based on body size, with 95% credible intervals shown as shaded ribbons. These are extracted from a model fit with an interaction between metadata and body size to purposefully investigate the influence of metadata and the relationship with body size. (B) Posterior distributions of the slope for body size (scaled and log10 transformed) under two model structures: V1 includes a random slope for body size by subrealm and a random intercept for metadata (as presented in main results), while V2 adds a random slope for body size by metadata. The inclusion of this additional random slope in V2 increases uncertainty and pulls slope estimates toward zero, particularly when data are sparse within body size measurement types as illustrated here. Also see Fig. S14.

**Table S1:**
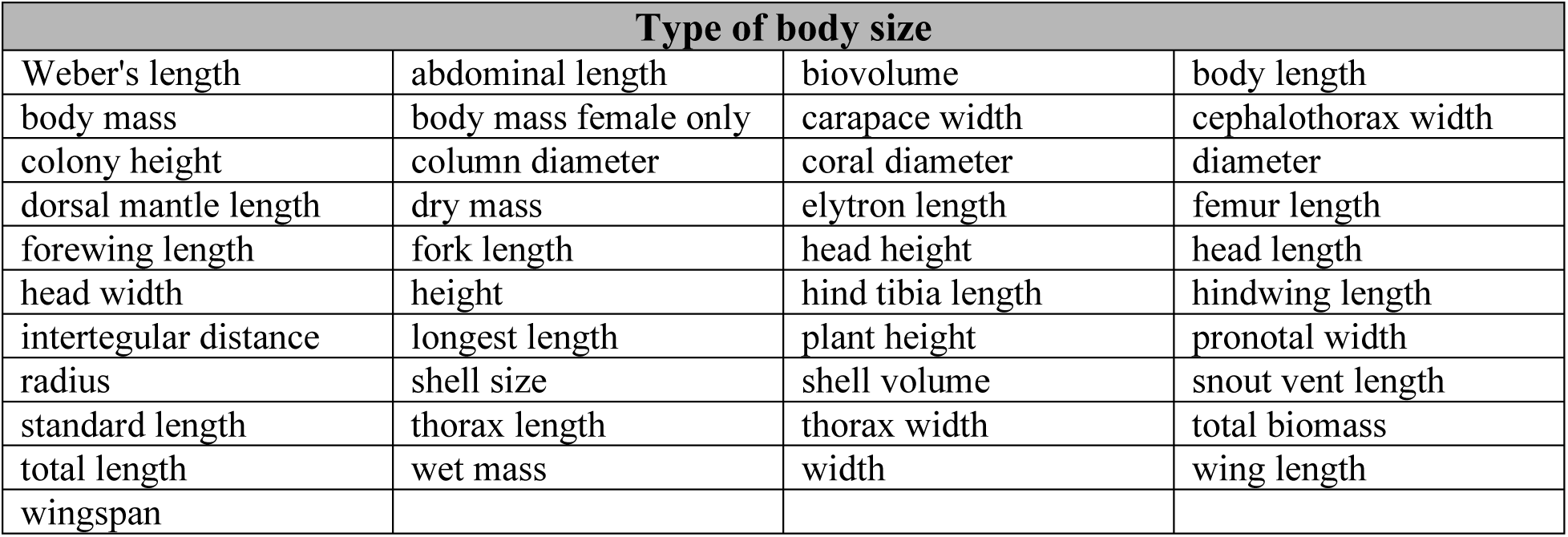
A list of 41 potential ‘types’ of body size that were used for potential inclusion in our body size dataset. We aimed to incorporate as many types of body size measures as possible and were not restrictive in our searching for body size measures.

**Data S1: (separate file)** The urban affinity values (N=56,181 total values (i.e., unique urban affinity score in a subrealm)) for potential inclusion in our analysis. This table is uploaded as a separate CSV file.

**Data S2: (separate file)** A final dataset for potential analysis and modeling, including a total of 94,087 observations (unique combination of species urban affinity values, subrealm, and body size measure) of 20,957 species that had at least one measure of urban affinity and at least one measure of body size. This table is uploaded as a separate CSV file. Note, however, that not every body size measure is made available due to a lack of permissions to share some datasets.

**Data S3: (separate file)** A list of the final potential ‘datasets’ that were used to aggregate measures of body size. Metadata refer to our naming scheme employed in our workflow; citation is a descriptor of either the paper citation or dataset citation or a descriptor of manual data aggregated by us; URL is the potential URL if applicable, and number of data points is the number of potential data points that could be used for the analysis. This table is uploaded as a separate CSV file.

